# MiniFAST: A sensitive and fast miniaturized microscope for *in vivo* neural recording

**DOI:** 10.1101/2020.11.03.367466

**Authors:** Jill Juneau, Guillaume Duret, Joshua P. Chu, Blake Madruga, Conor C. Dorian, Alexander V. Rodriguez, Savva Morozov, Daniel Aharoni, Jacob T. Robinson, François St-Pierre, Peyman Golshani, Caleb Kemere

## Abstract

Observing the activity of large populations of neurons *in vivo* is critical for understanding brain function and dysfunction. The use of fluorescent genetically-encoded calcium indicators (GECIs) in conjunction with miniaturized microscopes is an exciting emerging toolset for recording neural activity in unrestrained animals. Despite their potential, current miniaturized microscope designs are limited by using image sensors with low frame rates, sensitivity, and resolution. Beyond GECIs, there are many neuroscience applications which would benefit from the use of other emerging neural indicators, such as fluorescent genetically-encoded voltage indicators (GEVIs) that have faster temporal resolution to match neuron spiking, yet, require imaging at high speeds to properly sample the activity-dependent signals. We integrated an advanced CMOS image sensor into a popular open-source miniaturized microscope platform. MiniFAST is a fast and sensitive miniaturized microscope capable of 1080p video (1920×1080 pixels), 1.5 µm resolution, frame rates up to 500 Hz (achieved with windowing: 1920 × 55 pixels height) and high gain ability (up to 70 dB) to image in extremely low light conditions. We report results of ∼300 Hz *in vivo* imaging of freely behaving transgenic Thy1-GCaMP6f mice, high speed 500 Hz *in vitro* imaging of a GEVI and *in vivo* GEVI imaging in head-fixed mice. Our results extend miniaturized microscope capabilities in high-speed imaging, high sensitivity and increased resolution, opening the door for the open-source community to use fast and dim neural indicators.

## Main

Miniaturized microscopes used for *in vivo* neural recordings in small animals allow the activity of large populations of neurons to be visualized during the naturalistic behaviors that are critical to study how underlying neural circuits guide learning, behavior and decisions^1,2,3^. Development of miniaturized microscope technology has been rapid, with versions aimed at various applications such as wireless recording^4,5,6^, wide-field imaging^7^, optogenetic stimulation ^8^, volumetric imaging^9^, dual hemisphere imaging^10^, and audio triggered recording ^4^. Yet, limitations exist in frame rates, sensitivity, and resolution of the image sensors currently used.

Most miniaturized microscope technologies, use frames rates of 30 Hz or less, with the maximum reported being 60 Hz (Supplementary Table 1). Imaging genetically-encoded calcium indicators (GECIs), such as broadly used GCAMP6f versions, at frame rates of 30 Hz is adequate for many studies which are only concerned with slower Ca^2+^ dynamics. However, for applications such as transient detection^11^ or real-time processing, faster frame rates provide a distinct advantage in resolving the ∼40-ms rise times of the calcium transients ^12^ with high temporal precision and low latency. Moreover, in experiments aimed at studying fast network oscillations such as sharp wave ripples, ^13^ higher frame rates are necessary to capture faster frequency responses of the population dynamics. Using even faster neural indicators, such as genetically-encoded voltage indicators (GEVIs), would allow neural imaging experiments to resolve neural activity at the timescales of electrophysiology while retaining imaging advantages in non-invasiveness, genetic-specificity, and stability of interface. To take advantage of these benefits, miniaturized microscope frame rates will need to match these fast temporal kinetics (< 10 ms in some GEVI versions ^14,15^) requiring significant increases in frame rates (> 200 Hz).

Another important factor is the image sensor’s ability to image dim neural indicators. Due to the inherent tradeoff between exposure time and frame rate, imaging at faster frame rates results in reduced image brightness. When imaging fluorescent indicators such as GECIs and GEVIs, a higher signal can be achieved by increasing the excitation light power, but with the downside of increasing photobleaching of the fluorophores and potentially photodamaging tissue. The severity of photobleaching, linked to the excitation light power^13,16,17,18,19^, reduces the fluorescence signal over time and limits the experimental imaging time length. Thus, improvements in low-light imaging could allow for a reduction of excitation light power while still capturing images bright enough to detect optical activity. All other things being equal, the image sensor with maximum sensitivity, and to a lesser extent pre-digitization gain, will maximize performance.

While many miniaturized microscopes use a similar set of optics, most available miniaturized microscopes use image sensors with 0.5 megapixels and pixel sizes > 3 µm. While the resolution in these systems is constrained by the sensor pixel sizes rather than the optics, designers have largely settled on image sensors with larger pixel sizes to increase sensitivity. Yet, increasing the pixel density with smaller pixel sizes, would provide a greater number of samples between anatomical features and improve segmentation analysis to differentiate anatomical structure overlap.

Addressing the technology gaps stated above, in this study, we report MiniFAST, a fast and sensitive open-source miniaturized microscope module that provides imaging frame rates up to 500 Hz (achieved with windowing) and an excellent sensitivity/gain ability (up to 70 dB) to image in low signal environments. We show results using MiniFAST to image neural activity in freely-behaving GECI injected and transgenic mice, *in vitro* patch-clamp GEVI cells and GEVI injected head-fixed mice.

## Results

### MiniFAST design

To meet the design challenges of increased frame rates, sensitivity, and resolution, we designed a new open-source miniaturized microscope that integrates a Sony STARVIS back-sided illumination CMOS image sensor into our custom designed imaging PCB (Fig. 1, Supplementary Fig. 1, Supplementary Table 2). Compared to other miniaturized microscopes, MiniFAST’s image sensor has 8 times faster frame rates^3,4,8,9,20,21,22^ (Fig. 1c). While frame rates of 30 Hz or less have a full 1080p resolution (1920 × 1080 pixels), the image sensor can achieve faster frame rates by imaging subregions of the sensor (ex. 1920 × 55 pixels for 500 Hz). Moreover, the image sensor’s small pixel size of 2.9 µm provides increased spatial resolution when combined with optical lens sets typically used in miniaturized microscopes (Fig 1d).

**Figure 1:**
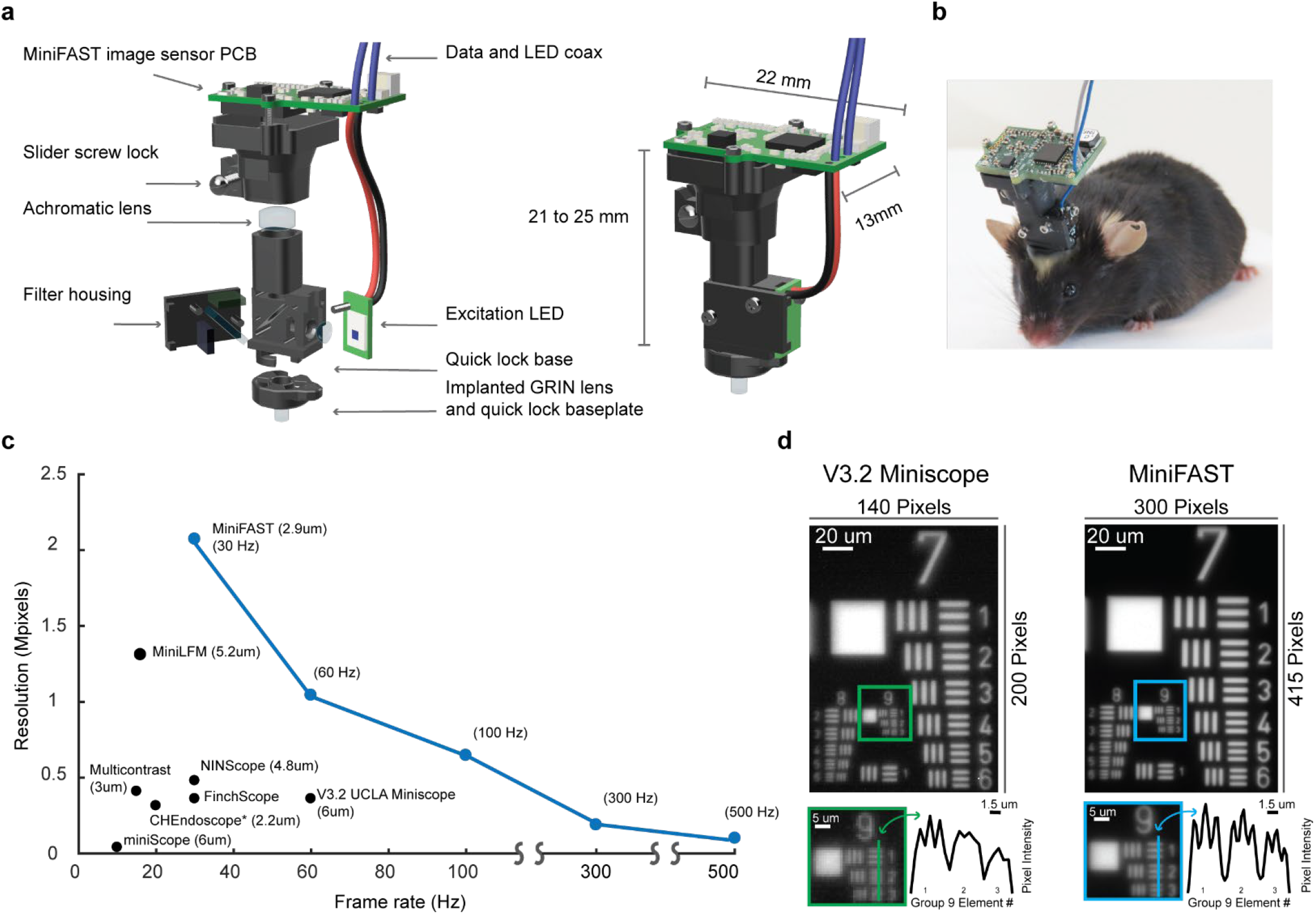
Published designs of miniaturized microscope image sensor resolution vs frame rate. **a**, Illustration of the MiniFAST miniaturized microscope imaging module. **b**, Image of a mouse wearing the microscope. **c**, Figure of a comparison of miniaturized microscope image sensor resolution versus published frame rates^3,4,8,9,20,21,22,23^. Numbers in parentheses indicate the sensor pixel size. *Note: CHEndoscope image sensor is 5 MP, but images are binned to 648 × 486 to achieve a 20 Hz frame rate. **d**, Images from V3.2 Miniscope (left) and MiniFAST (right) of a USAF 1951 high resolution test target cropped to show groups 7 through 9. V3.2 Miniscope settings are 10 Hz, 0 dB gain, 0.18 mW excitation light. MiniFAST settings are 8 Hz, 0 dB gain, and 0.1 mW excitation light. The same set of microscope housing, filters and optical lenses was used on both microscopes. Inset shown is of the line profile through Group 9 elements.

MiniFAST’s image sensor PCB and optical housing (Fig. 1a-b) builds on the V3.2 UCLA Miniscope^3^, a popular open-source miniaturized miniscope platform, and is compatible with its existing data acquisition module and user GUI. Although MiniFAST’s image sensor PCB can still mate to the V3.2 Miniscope housing, we also designed a custom 3-D printed MiniFAST housing and baseplate which uses keyed locks and magnets to create a quick mating connection. We have found this design significantly reduces animal handling time compared to the V3.2 Miniscope attachment method. Other aspects of MiniFAST’s optical housing are the same as the V3.2 Miniscope incorporating the same optical lens/filter set and manual focus slider. The weight of the MiniFAST module including the image sensor PCB, optical housing, optics, and LED is 3.45 grams and is comparable to the V3.2 Miniscope’s weight of 3.2 grams.

To compare the resolutions of the V3.2 Miniscope and MiniFAST, a resolution test was performed using a USAF 1951 high resolution test target (Edmund Optics #55622) (Fig. 1d). Using the same set of optics, similar frame rates and gain settings, MiniFAST (2.9 µm/pixel) resolved line pairs down to Group 9, Element 3 (645.1 line pairs/mm) and showed an increase in contrast across all Group 8 and Group 9 elements when compared to the V3.2 Miniscope (6 µm/pixel). Binning with a 2×2 average format of the image taken with MiniFAST reveals a similar resolution to the V3.2 Miniscope, demonstrating the increased resolution is achieved by MiniFAST’s decreased pixel size (Supplementary Fig. 2).

### *In vivo* – Imaging GECI place cell activity in freely-behaving GECI injected mice

To evaluate neural imaging in live mice, after injection of adeno-associated viruses (AAV), we imaged 2 mice transiently expressing the calcium indicator GCaMP6f^12^ in excitatory neurons in the CA1 region of the hippocampus (Supplementary Video 1 and 2). In one of the mice, during a 17-minute exploration session, we imaged over 100 neurons and detected the expected spatial tuning (Fig. 2, Supplementary Fig. 3). Our novel design was stable over a 2-hour exploratory imaging session (Supplementary Fig. 4) and over month-long periods (Supplementary Figs. 5 and 6) verifying the integrity of the quick-lock baseplate and surgical attachment methodology.

**Figure 2:**
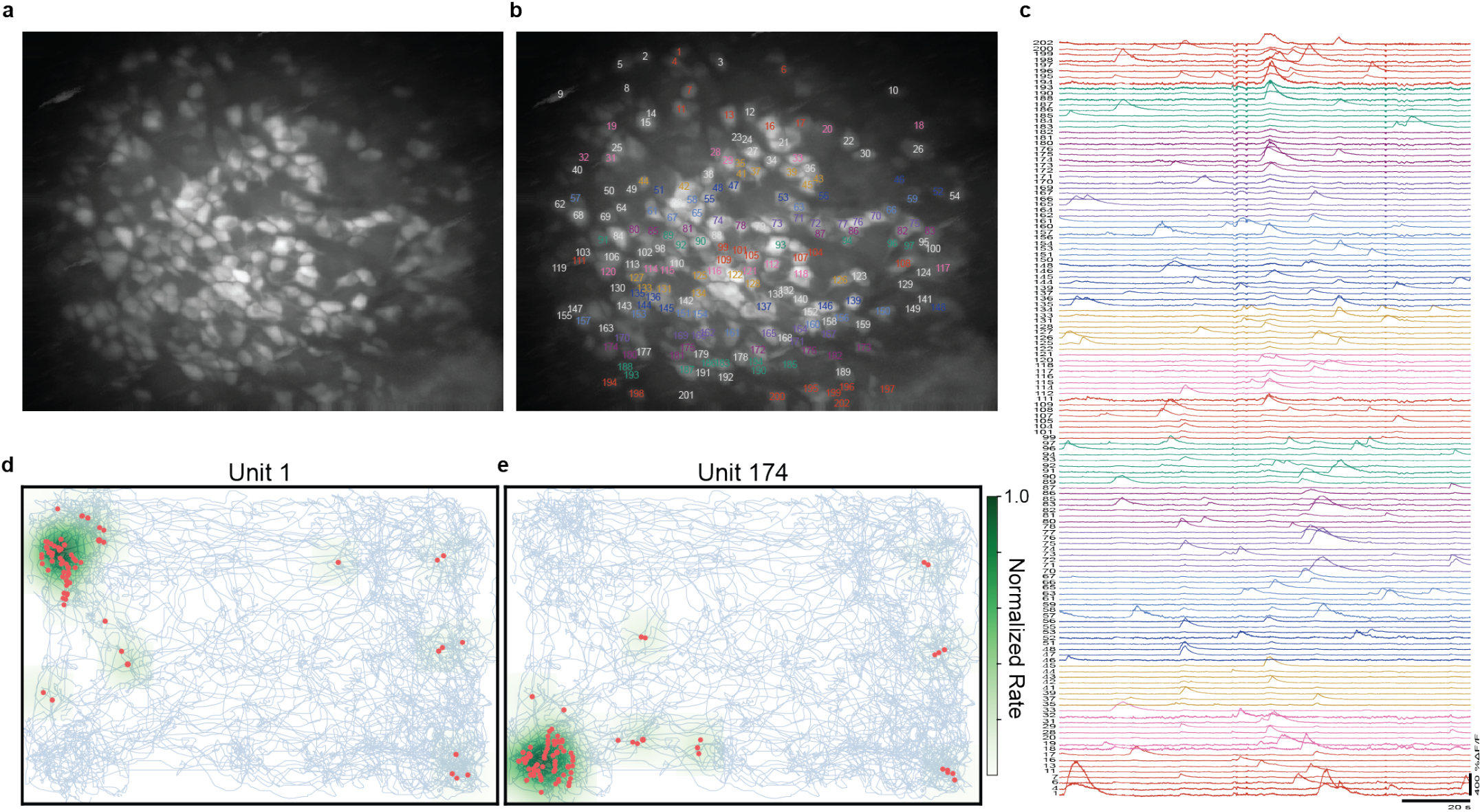
*in vivo* CA1 place cell imaging of AAV GCaMP6f injected mouse. **a**, Maximum intensity projection images (1320 × 1040 pixels) from Ca^2+^ imaging in the CA1 region of hippocampus of a mouse exploring an open field. MiniFAST settings were 30 Hz, 15 dB gain, and 0.4 mW excitation light. **b**, (**a**) with overlaid neurons labels. **c**, Calcium activity traces for neurons with detected transients during a 120 second period. Numeric labels match **(b)**. Scale bars are 20 s and 400 ΔF/F. **d, e** The spatial tuning for two example cells shows the expected place-field. Spike events (red) and position trajectories (blue) observed during exploration of a 60 × 40 cm open field are overlaid on the inferred rate map (green).

### *In vivo* – Imaging GECI activity in freely-behaving GECI transgenic mice

Transgenic Thy1-GCaMP6f GECI mice are broadly used, providing stable expression of GECIs in pyramidal neurons over long time scales without the necessity of viral injections^24^. One potential drawback is that neurons in the Thy1-GCaMP mice often exhibit dimmer and faster fluorescence signals compared to those generated using viral transfection^13,24^. To further assess MiniFAST’s ability to resolve activity of dim neural indicators, we imaged CA1 neurons of a Thy1-GCaMP6f GP5.17 mouse exploring an open field at frame rates of 30, 60, 100, 200 and 293 Hz (Fig. 3). Per the image sensor’s specification, faster frame rates were achieved by adjusting the window row size. Next, the window was positioned over the same cell for the 30, 60, 100, and 200 Hz sessions. However, for the 293 Hz traces, a nearby region with more active cells was selected, revealing three distinct cells in the field-of-view (FOV). Despite the dim signal, MiniFAST resolved Ca^2+^ activity in the cells at each frame rate, even at the challenging 293 Hz frame rate (Fig. 3a-5, Supplementary Fig. 7).

**Figure 3:**
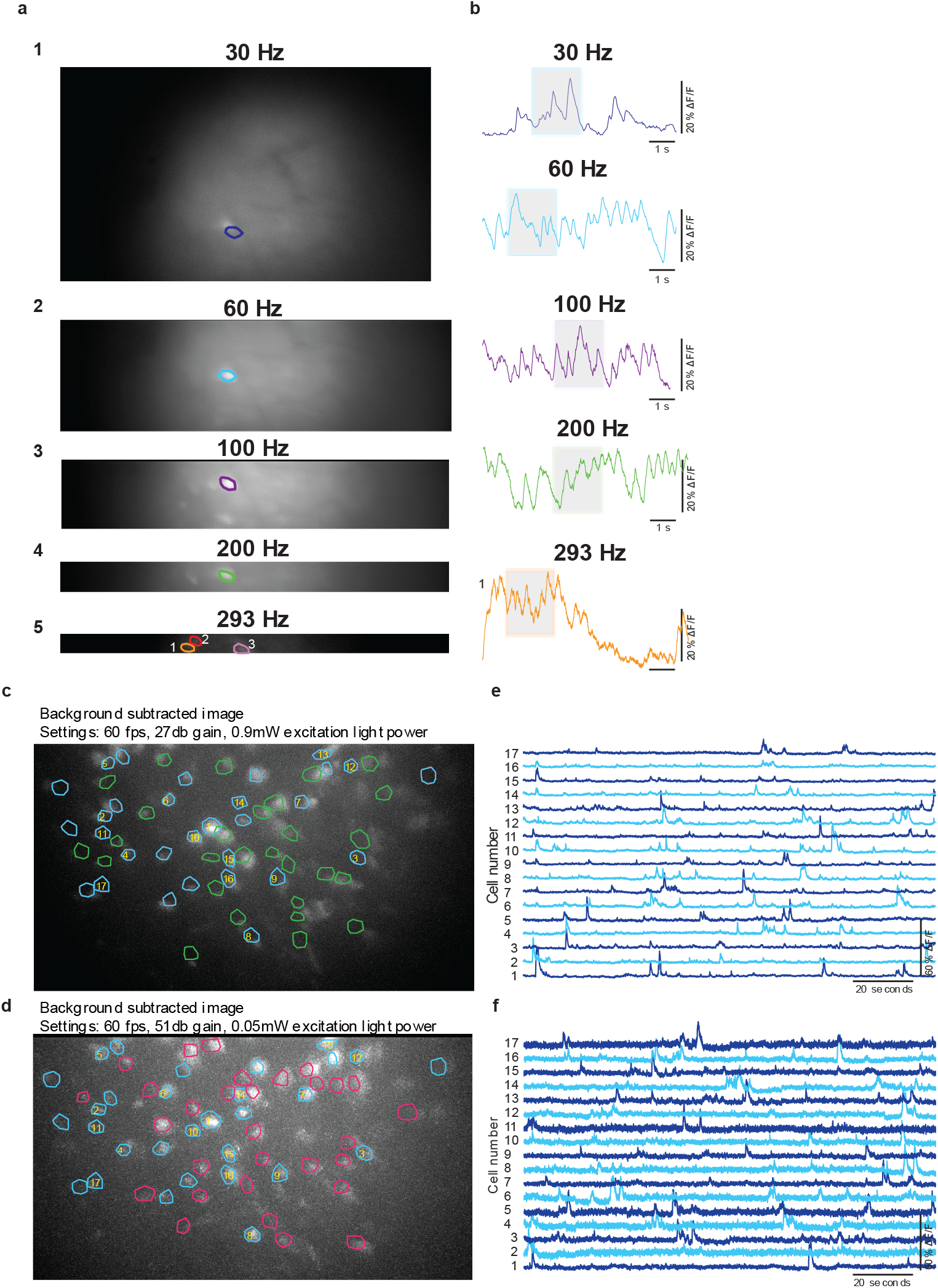
*in vivo* imaging of a freely-behaving GECI transgenic mouse (30 – 293 Hz) **a**, Raw maximum projection image from imaging sessions of the same mouse at 30 Hz (1800 × 1040 pixels), 27 dB gain, 0.86 mW excitation light power (**a1**), 60 Hz (1900 × 544 pixels), 27 dB gain, 1.4 mW excitation light power (**a2**), 100 Hz (1900 × 337 pixels), 27 dB gain, 2.5 mW excitation light power (**a3**), 200 Hz (1900 × 156 pixels), 30 dB gain, 2.5 mW excitation light power (**a4**) and 293 Hz (1900 × 102 pixels), 30 dB gain, 3.25 mW excitation light power (**a5**). **b**, corresponding 8-second long Ca^2+^ traces of one cell in each FOV. **c-d**, Background subtracted images and circled ROIs for 5-minute imaging sessions. **c**, Excitation light power 0.9 mW, **d**, Excitation light power 0.05 mW. Blue outlined ROIS are ROIs active in both sessions. Green and magenta outlined ROIs are regions active only in that specific session. **e, f**, Ca^2+^ activity traces for blue outlined cells with yellow text labeled in (**c**) and (**d**) that were active over the 120 second period in the 0.05 mW session shown in (**d**).

While slower dynamics of the Ca^2+^ activity were captured at frames rates as low as 30 Hz, imaging at higher frame rates revealed faster Ca^2+^ signal kinetics, offering a greater number of data samples for the onset of Ca^2+^ transient edge detection^34,35,36^. This is particularly crucial in experimental paradigms employing real-time closed-loop systems. For example, in a Ca^2+^ edge occurring over a 150ms interval, 30 Hz imaging yields only 5 samples compared to 43 samples when imaging the same cell at 293Hz (Supplementary Fig. 7). Using slower 30 Hz full-frame imaging proves beneficial for identifying specific cell activity, such as place cells, within a larger subset of cells. Subsequently, employing faster frame rates with windowing enables a closed-loop system with rapid detection and response to a specific cell’s Ca^2+^ edges.

Leveraging the wide range of gain settings available on MiniFAST’s image sensor, we wanted to evaluate MiniFAST’s ability to detect Ca^2+^ activity at ultra-low excitation light levels where photobleaching effects could be minimized. In a freely behaving Thy1-GCaMP6f GP5.17 mouse exploring an open field, we imaged the same cell region in two 20-minute sessions (60 Hz) each with different excitation light/gain levels, to determine if activity of the same cells could be consistently detected in both sessions (Supplementary Video 3). The settings were chosen so that in the higher excitation light session (0.9mW/27db gain) the cell outlines could clearly be seen in the 60 Hz frame video compared to very little cell contrast in the low excitation light session (0.05mW/51 db gain). Analysis of activity from the 0.9 mW excitation light session (58 cells) (Fig. 3 c, e) and a low 0.05 mW excitation light session (60 cells) (Fig. 3d, f) revealed 30 cell ROIs active in both sessions.

Importantly, analysis of the Ca^2+^ activity demonstrated calcium spiking events were still clearly detectable in the low 0.05 mW excitation light power imaging sessions (Fig. 3f). These results extend the previous demonstration of the 2-hour imaging sessions of 2 injected AAV GCaMP6f mice (Supplementary Fig. 4) where Ca^2+^ activity was detected throughout using excitation light settings of 0.2mW and 0.05mW excitation light, but with imaging at only 30 Hz frame rates.

### *In vitro* – Imaging GEVI cell patch clamp

Next, in order to test MiniFAST’s ability to image fast and dim neural indicators such as genetically-encoded voltage indicators (GEVIs), we imaged patch clamped HEK293T cells transfected with the GEVI ACE2N-mNeon^15^ (Fig. 4a). During 500 Hz imaging sessions with MiniFAST, using a voltage-clamp mode, we electrically evoked a voltage step of -70 mV to +40 mV for pulse widths of 2 and 5-ms at 10 and 50 Hz to produce fluorescence transients. A comparison of electrical recordings with optical recordings revealed that MiniFAST reproduced the induced pulse trains (Fig. 4c-h). Importantly, even at stimulation pulse widths of 2-ms at 50 Hz, each pulse was still detectable with an average peak ΔF/F of -3.8% (Fig. 4h).

**Figure 4:**
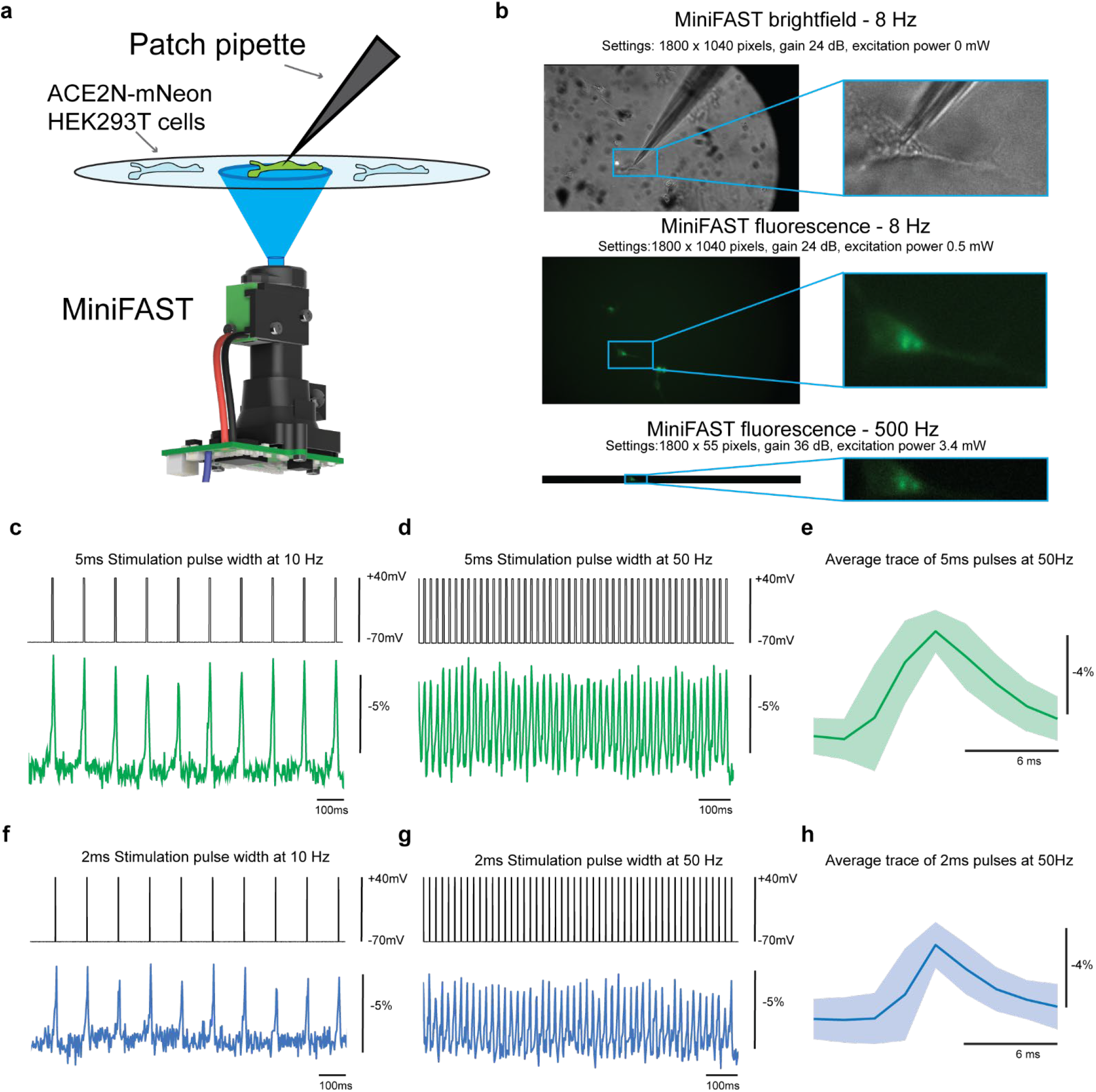
*in vitro* high-speed imaging (500 Hz) of a genetically-encoded voltage indicator. **a**, Illustration of the setup used for imaging and patch clamp. **b (top)**, Full-frame brightfield image and blue-framed inset of zoomed image of the exact cell used for the data shown in **(c-h), b (middle)**, False-colored fluorescence image of full frame at 8 Hz. **b (bottom)**, False-colored fluorescence image of reduced frame at 500 Hz. Per image sensor specifications, faster frame rates were achieved by decreasing window sizes. **c-d, f-g**, Raw traces from patch clamp recordings of the GEVI ACE2N-mNeon transfected in HEK293T cells. Electrical recordings (top) and MiniFAST optical recordings (bottom) over 1-second long pulse trains with steps of -70 mV to +40 mV. **c-d**, 5-ms Pulse width stimulation at 10 Hz (**c**) and 50 Hz (**d**). **f-g**, 2-ms Pulse width stimulation at 10 Hz (**f**) and 50 Hz (**g**). **e, h**, Average peak-aligned traces, and background shaded with 2.5% and 97.5% empirical values for the 50 Hz pulse traces. Average peak ΔF/F for 50 Hz of 5-ms was -5.4%. Average peak ΔF/F for 50 Hz of 2-ms was -3.8%. **c-h** MiniFAST settings were 500 Hz, 36 dB gain, 3.4 mW excitation light and a window size of 1800 × 55 pixels.

### *In vivo* – Imaging GEVI activity in head-fixed mice

To demonstrate the ability of MiniFAST’s image sensor (Sony IMX290LLR) to measure spiking of GEVI cells *in vivo*, an IMX290 development board was used as the CMOS camera in a custom-built single-photon epi-fluorescent microscope with the setup described previously^34^. We imaged ASAP3-Kv in dorsal CA1 SST^+^ interneurons through a deep window while mice were awake, head-fixed and running. Since the image sensor has a reduced window at high frame rates, the window of 1920×47 pixels was placed over the cell in the FOV. Blue 455nm light-emitting diode (LED), illumination at 25mW mm^-2^, revealed spikes (cell 1: 64 spikes in 4.5s, cell 2: 34 spikes in 5 s), including resolving ensembles of spikes, while imaging at 500 fps (Fig 5 a,b) with average spike df/f response of ∼4.2% and (Fig 5 e). NOSA^32^, an open-source analytical toolbox for multicellular optical electrophysiology designed specifically for the challenges of GEVI data analysis, was used to detect spikes by using a conservative spike threshold setting of 4.5 standard deviations of noise (Supplementary Fig. 9).

**Figure 5:**
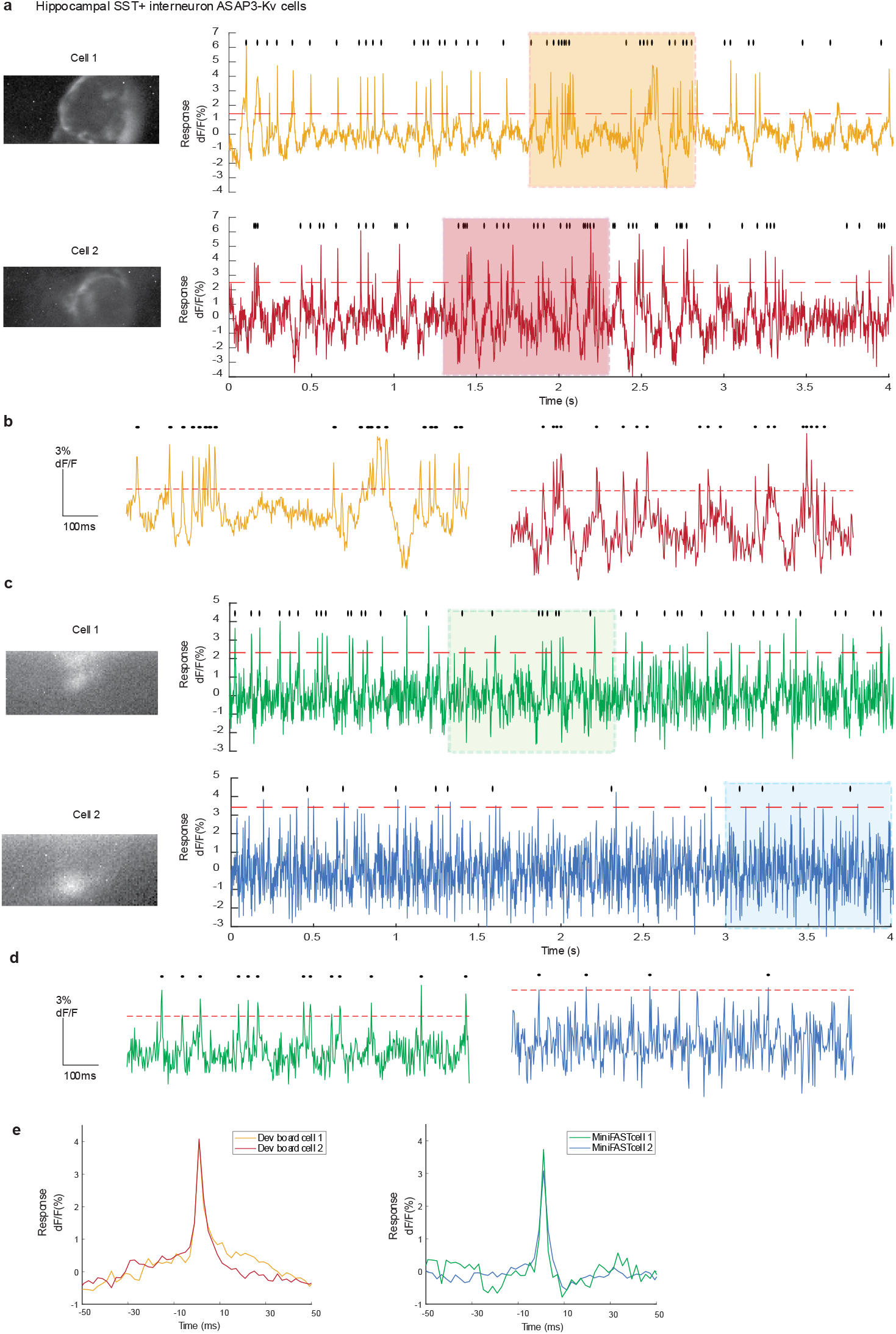
*in vivo* high-speed imaging (500 Hz) of ASAP GEVIs in hippocampal SST^+^ interneurons in head-fixed mice. **a**, Example images and GEVI traces from in vivo interneuron ASAP3-Kv cells imaged with a Sony IMX290 development board as the CMOS camera in a custom-built single-photon epi-fluorescent microscope imaging an awake, head-fixed running mouse. The cell images are IMAGEJ standard deviation z-projection of the ROI imaged at 500fps. ASAP3-Kv fluorescent traces were acquired at 500fps, camera gain of 30-40db, using light excitation power of 25mW mm^-2^. Raw traces are background subtracted. ASAP3-Kv is a high to low indicator and the traces shown here are inverted. The red dashed line represents a spike threshold of 4.5 standard deviations of the noise and black dots represent detected spikes. **b**, 1-second expanded traces from (**a). c**,**d**, Example images and GEVI traces from in vivo interneuron ASAP3-Kv cells imaged with MiniFAST microscope secured in head-fixed setup in the same mouse as **(a**,**b)**. Given the setup differences, we could not verify the same exact cells for both setups. The images and GEVI traces were processed in the same way as in **(a**,**b)**. ASAP3-Kv fluorescent traces were acquired at 500fps, camera gain of 37db, using light excitation power of 10mW mm^-2^. **e**, Average spike event shape for the 2 cells shown for the IMX290 development board setup and the MiniFAST setup.

After verification the Sony IMX290 image sensor could detect GEVI spikes, we replaced the bench microscope setup with MiniFAST (Sony IMX290 image sensor PCB, optical housing, attached grin lens and slightly modified LED PCB), in a fixed position over the deep window of the same awake, head-fixed, running mouse. We imaged cells with MiniFAST at 500fps with blue 470nm LED illumination. While an increased noise level was observed, cells were clearly identified and neural activity could be detected (Fig 5 c, d). Using NOSA, with the same settings as before (spike threshold set to 4.5 standard deviations above the noise), 74 spikes were detected in cell 1 over an 8 s imaging period and 25 spikes were detected in cell 2 over a 10 s imaging period with average spike df/f response of ∼3.5-4% (Fig 5 e, Supplementary Fig. 10-13). To investigate if the cells were spiking individually or due to activity in the surrounding cells, an ROI off the cell, but nearby, was selected and using the same threshold as the on-cell region, only 8 spikes were detected in cell 1 and 5 spikes in cell 2 over the same imaging periods (Supplementary Fig. 10-12). These results indicate that miniaturized microscopes can detect GEVI activity in vivo.

## Discussion

Overall, MiniFAST was designed to provide the open-source research community a miniaturized microscope capable of high-speed imaging (up to 500 Hz), high sensitivity to image in dim conditions (gain up to 70 dB) and increased resolution (1920 × 1080 pixels and a 2.9 µm pixel size). To our knowledge, we are the first to demonstrate *in vivo* imaging using a miniaturized microscope at frame rates of ∼300 Hz while resolving Ca^2+^ activity with dim signals from a Thy1-GCaMP6f transgenic mouse exploring an open field (Fig. 3). Furthermore, also with Thy1-GCaMP6f mice exploring an open field, we have shown the potential of using ultra-low excitation power settings (0.05 mW) to reduce the severity of photobleaching while still resolving Ca^2+^ activity at 60 Hz frame rates (Fig. 3). Additionally, we have been the first to demonstrate use of a miniaturized microscope to image GEVI kinetics at frame rates of 500 Hz, resolving 2 ms duration voltage pulses at 50 Hz *in vitro* (Fig. 4) and resolving GEVI spikes *in vivo* in head-fixed mice (Fig. 5).

Furthermore, this work demonstrates that inexpensive rolling shutter image sensors, over the traditionally used global shutter sensors, can successfully image GEVIs at high speeds. Typically, rolling shutter image sensors suffer when imaging objects at high speeds where image distortions can occur if the objects are moving across the FOV at speeds faster than the imaging frame rates. In the context of imaging neural activity, where cells are relatively stationary despite rapid changes in fluorescence dynamics (< 10ms), these distortions are mitigated. However, it is important to note, with rolling shutters, each row is exposed at a slight offset to the next row. In the case of 500 Hz frame rates with a frame time of 2ms, each row experiences an offset by ∼40us. Consequently, there will be an exposure time lag vertically across a cell depending on how large the cell is in the FOV. Despite these inherent characteristics, as evidenced by the patch clamp data (Fig. 4), MiniFAST successfully resolved the GEVI activity given the ∼6ms fluorescence change. If imaging a faster GEVI, the rolling shutter time lag issues would likely become more pronounced.

While these results demonstrate that a small CMOS image sensor can image and detect GEVI spikes *in vivo*, further work is required to use miniaturized microscopes practically for freely behaving GEVI imaging in mice. First, as demonstrated by the higher noise floor seen in the MiniFAST setup compared to the bench microscope setup, a more directed excitation light path would improve the SNR significantly. Currently, the MiniFAST LED optics generate light that spreads over the entire FOV, exciting all cells in the region rather than “spot-lighting” individual cells. Also, given the sensitivity of GEVIs to photobleaching, a more directed light path and automation of the light direction for shorter durations could reduce photobleaching enabling longer imaging sessions. Second, the current MiniFAST/UCLA DAQ uses the IMX290’s parallel data interface. If the system was redesigned to use the image sensor’s CSI/MIPI interface, a larger window can be used at higher speeds. For example, at 500 fps, the row size for the current MiniFAST setup is ∼50 pixels vs ∼250 pixels if the CSI interface was used (Supplementary Fig. 14). This would enable high speed imaging in multiple cells at once and combined with automated windowing and a directed light path, practically imaging GEVIs in freely-behaving mice is feasible. Third, Sony has released newer versions of the STARVIS series 2Mpixel image sensors such as the IMX462 and IMX662 which are supposed to provide even greater low light sensitivity.

Another important consideration in using miniaturized microscopes with larger pixel counts or imaging at high speeds is the increased data size that will be accumulated. For example, a researcher using MiniFAST for standard GECI imaging at the typical 30 Hz must consider if the smaller pixels/larger pixel count is worth 4 times the amount of imaging data (2GByte for every 30 seconds of 30 Hz data) that will need to be processed. Also, many of the open-sourced algorithms such as those used for motion correction, automatic ROI and spike detection were designed for smaller datasets and using the same code for a 2Mpixel image sensor requires more hard drive storage and processing time/power. Therefore, if an experiment calls for standard GECI imaging, a researcher might opt for one of the other open-source microscopes that are more suited for smaller datasets. However, for 30Hz imaging, MiniFAST can be advantageous towards experiments that would benefit from the ability to image anatomical structures smaller than neurons or wide-field imaging where single-cell neurons might be distinguished with small pixel pitches. Additionally, as shown here, MiniFAST can be used at slower frame rates for longer experiments that require very low light levels to prevent photobleaching or potentially for emerging neural indicators such as bioluminescent indicators which are promising because they eliminate the problems of autofluorescence and photobleaching that occurs in fluorescent indicators but are significantly dimmer than GECIS in current versions^25,26^. Post processing down-sampling of the datasets can also help to reduce the amount of data that would need to be stored.

In conclusion, while this paper demonstrates advancements to bridge the technology gap, we expect advancements in image sensor technology and protein engineering will continue to provide new capabilities in miniaturized microscopes.

## Methods

### MiniFAST imaging sensor module and software

MiniFAST is based on a previously described popular open-source miniscope platform, V3.2 Miniscope^3^, and consists of a custom designed image sensor PCB, a modified optical housing and microscope baseplate. With updated software, MiniFAST’s image sensor PCB is compatible with the existing miniscope data acquisition board (DAQ) and Window’s based GUI. MiniFAST’s CMOS image sensor (Sony IMX290LLR), was chosen for its ability to achieve the design goals of increased resolution, frame rates and sensitivity. As in the previous design, the image sensor PCB design incorporates a single coax cable for power, image sensor data and commanding as well as one additional small wire to externally power the onboard light excitation LED. Powering the LED externally reduces excitation light flicker due to the increased current draw of the image sensor. The external power supply of 4 V requires a small modification to the DAQ board. MiniFAST’s image sensor PCB weighs 2 g compared to 1.7 g for the image sensor PCB of the original system.

The MiniFAST optical housing is similar in design to the V3.2 Miniscope’s optical housing and both incorporate the same achromatic lens, filter sets, excitation LED (LUXEON Rebel Color Blue, 470 nm), half ball lens and GRIN Lens. As before, the focus depth is adjustable by manually sliding the microscope housing. However, MiniFAST’s optical housing is 3-D printed and we have found using the Formlabs Form2 black resin to be the least auto-fluorescent in our imaging wavelengths as previously reported^4^. MiniFAST also incorporates a larger base to accommodate the larger image sensor size and includes a quick lock baseplate. Importantly, the microscope quick lock base, also 3-D printed, does not require any additional set screws to mate to the optical housing. The mating of the optical housing and the quick lock base uses keyed locks and magnets to hold the microscope firmly in place. This new design results in reduced animal handling time and the need for animals to be anesthetized when following handling techniques previously set forth by Gouveia et al.^27^. Moreover, MiniFAST’s optical housing, lens set, filter set, LED and baseplate weighs 1.6 g compared to 1.95 g for the original system. The set of filters used during the imaging experiments was the following: excitation filter (Chroma ET470/40x), dichroic mirror (T495lpxr), and emission filter (ET525/50m). Also, the optical set included a half-ball lens (Edmund Optics #47-269), one achromatic lens from the following set (10mm - Edmund Optics #45-207, 15mm - Edmund Optics #45-207, 20mm - Edmund Optics #45-208), and a GRIN lens (Edmunds Optics part numbers noted in text).

The V3.2 Miniscope DAQ firmware and Windows GUI have been updated to integrate the new image sensor’s features such as increased gain, frame rate, black offset, and window position. As before, the GUI supports the ability to simultaneously sync frames of both the microscope and a behavior camera as well as to create a synced 1-ms resolution timestamp file. Notably, during the patch clamp *in vitro* testing with frame rates up to 500 Hz, the timestamp file accurately aligned the video frames to the timing of the patch clamp voltage pulses.

### In vitro whole-cell patch clamp

#### MiniFAST setup

A 0.2 mm working distance (WD) GRIN lens (1.8 mm diameter, 4 mm length, Edmund Optics part #64-520) was secured to a MiniFAST baseplate with superglue and attached to a MiniFAST microscope. The microscope was then connected to a micromanipulator with an in-house designed 3-D printed bracket. The microscope and micromanipulator setup were positioned so that the GRIN lens of the microscope was directly under an *in vitro* cell recording chamber. For each patch clamp session, cells were loaded into the recording chamber. The microscope was then focused onto the cells through fine position control using the micromanipulator. An external lamp provided the light source for the brightfield images. MiniFAST’s onboard LED provided the excitation light during fluorescent imaging. To initially find the cells in the field-of-view (FOV), MiniFAST was set to use a full frame (1920 × 1080 pixels), a low frame rate of 10 Hz and an excitation light of ∼1 mW. After a cell was patched, the microscope settings were changed to the decreased window size required by the faster frame rate (1800 × 55 pixels @500 Hz) and the window was positioned to include the cell in the FOV. Then, while pulse voltage trains were delivered through the patch clamp, the cell was imaged at 500 Hz, gain 36 dB, and excitation light of 3.4 mW.

#### Cell culture preparation

The plasmid ACE2N-mNeon^15^ was a gift from the Francois St. Pierre lab. HEK293T cells obtained from ATTC were replated on 24 well plates prior to transfection. The cells were transfected using Lipofectamine 2000 Transfection Reagent (Invitrogen) following the manufacturer’s recommendations. Briefly, for each well, 1 ug of plasmid DNA was combined with 1 uL of Lipofectamine 2000 in 50 uL OptiMEM, incubated at room temperature for 20 minutes, and added to 70% confluent HEK293T cells. The plasmid bearing the coding sequence for ACE2N-mNeon under a CMV promoter were purified using ZymoPURE™ II Plasmid Midiprep Kit.

#### Electrophysiology

The cells were held in a recording chamber filled with extracellular recording buffer (in mM: 145 NaCl, 5 KCl, 3 MgCl2, 10 HEPES and 1 CaCl2; pH 7.2; adjusted to 320 mOsm with sucrose). Glass patch pipettes (resistance of 3 to 5 MΩ) were filled with intracellular buffer (in mM: 140 KCl, 10 HEPES and 0.5 EGTA; pH 7.2; adjusted to 320 mOsm with sucrose) and contacted the cell membrane to generate seals ≥ 1 GΩ. For the whole cell configuration, a negative pressure of -70 mmHg was applied inside the pipettes until the cell membrane ruptured. An Axopatch 700 A recorded currents under voltage clamp conditions. The recorded current was filtered at 10 kHz and digitized at 2 kHz using a Digidata 1550 (Molecular Device). For voltage pulses, the cell’s holding potential was set at -70 mV and increased to +40 mV with 2-ms and 5-ms pulse durations at frequencies of 10 Hz and 50 Hz.

#### Animals

All experimental protocols were approved by the Institutional Care and Use Committee of Rice University and adhered to the National Institute of Health guidelines. For *in vivo* studies using GECIs, we used male and female mice that were either wild type (WT) C57BL/6 from Charles River or transgenic C57BL/6J-Tg (Thy1-GCaMP6f) GP5.17Dkim/J from Jackson Laboratories, age > 5 weeks and weight > 20 grams. Mice were group housed until the lens implant surgery after which they were individually housed.

For *in vivo* studies using GEVIs, all experimental protocols were approved by the Chancellor’s Animal Research Committee of the University of California, Los Angeles, in accordance with the National Institutes of Health (NIH) guidelines. We used male mice that were of type SST-IRES-Cre (https://www.jax.org/strain/013044). The mice were 35 weeks at surgery and 39 weeks at experiment time.

#### Viral construct

AAV9.CamKII.GCaMP6f.WPRE.SV40 (Titer≥ 1×10^13^ vg/mL) was purchased from Penn Vector Core.

### GECI AAV injection, lens implant and microscope baseplate surgeries

For *in vivo* imaging experiments, mice underwent stereotaxic surgeries consisting of an AAV injection (WT only mice), lens implant and a baseplate implant. The surgical techniques align with methods described in^3,5^, but with changes in bonding adhesives. To summarize, for AAV injection and lens implant surgeries, surgery preparation consisted of sterilizing all tools and surgery bench area using autoclave cycles or chemical methods. The GRIN lens was sterilized in a 12-hour ethylene oxide cycle. For all surgeries, mice were anesthetized with 1.5-2.5% isoflurane and 1L/min oxygen. The animal was then placed in a stereotaxic apparatus with the animal resting on a water heater pad and draped with Press’n Seal Cling Film^28^ to maintain body temperature and provide a sterile field. Ophthalmic ointment was applied to the eyes. The incision site was cleaned with a chlorhexidine scrub, and Lidocaine was administered below the incision site. Throughout the surgery, saline injections were administered subcutaneously (SC) to maintain hydration levels. For the AAV and lens surgeries, mice were SC administered Buprenorphine Slow Release (1 mg kg^-1^) for post-operative pain and Dexamethasone (4 mg kg^-1^) to reduce brain edema.

In the WT mice for the AAV injection surgery, we unilaterally injected 500nl of AAV9.CamKII.GCaMP6f.WPRE.SV40 (Penn Vector Core / Addgene) into the dorsal CA1 region of the hippocampus (−2mm anterior-posterior (AP), +2 mm medio-lateral(ML) and -1.65mm ventral positioned relative to bregma) using a Hamilton syringe with an injection rate of 50 nl/min. One week after the AAV injection for the WT mice or as a first surgery on the transgenic mice, a GRIN lens implantation surgery was performed. After the incision was made and skull surface cleaned, the craniotomy location was marked, and the skull was lightly scored with a scalpel. A thin layer of C & B Metabond Cement (Parkell) was then applied to exposed skull areas except at the site of the craniotomy. After waiting five minutes for the Metabond to cure, a 2 mm craniotomy was performed with the center at location -2 mm AP and +1.5 mm ML. Cortical tissue was slowly aspirated using a 30-gauge blunt needle while continually applying sterile saline. Aspiration ceased once the second layer of the corpus callosum striations appeared. After aspiration was complete, bleeding was reduced by placing a small section of gel foamed soaked with sterile saline over the craniotomy site for five minutes. An in-house designed custom 3-D printed lens holder and vacuum line was attached to a stereotaxic bar. A GRIN Lens (1.8 mm diameter, 4 mm length, Edmund Optics #64-519), vacuum secured with the lens holder, was lowered to 1.35 mm below the surface of the skull. With the lens still vacuum secured in the lens holder, a thin layer of Flow-It ALC Dental Composite (Pentron) was applied around the interface of the skull and GRIN lens and then immediately cured for 20 seconds using a UV Curing light (NKSI LY-A180). Application of small amounts of Flow-It and curing continued until a cured layer secured the lens to the skull surface.

Then, the vacuum was turned off and the lens holder was slowly lifted away from the lens. To protect the lens, Kwik-Sil (WPI) was added on top of the lens and allowed to cure for 10 minutes. At the end of both the AAV injection surgery and lens surgery, Meloxicam (2 mg kg^-1^) was administered SC. Meloxicam, Dexamethasone and saline injections were post-operatively given the two days following surgery.

Three weeks after the lens implant surgery, a baseplate surgery was performed. Before the surgery, the sides and bottom of a 3-D printed baseplate was lightly scored to ensure adhesive bonding. During the surgery, with the mouse under anesthesia and secured in a stereotaxic apparatus, the Kwik-Sil was carefully removed from the skull and lens area. Then, the baseplate was secured to a MiniFAST microscope and placed over the lens.

With the microscope powered on, the best FOV was found by manually changing angles of the microscope. With the microscope held in the desired position, small amounts of Flow-IT were slowly applied and UV cured to areas below and around the baseplate. Once the microscope and baseplate were completely secure, the microscope was removed from the baseplate and a protective lens cap was then placed over the baseplate.

Meloxicam and saline injections were administered post-operatively. Imaging sessions occurred 3-5 days after the completion of the baseplate surgery.

### *GECI In vivo* imaging sessions

After recovering from the baseplate surgery, the mice were habituated to handling^27^ and to carrying a sham microscope. During imaging sessions, with a MiniFAST microscope attached to the mouse’s baseplate, the mice were placed in a 65 cm by 40 cm open field and allowed to explore freely. A 1.5m 36-gauge data coax cable was used for the main microscope cable and a 2.5m M17/113-RG316 cable was used as an extension to the DAQ. The upgraded Miniscope Windows GUI was used to control and record frames from MiniFAST and sync a behavior camera (Logitech C920 HD Pro Webcam) at 30 Hz.

For the photobleaching experiments, the mice explored the open field for 10 minutes with the microscope attached but powered off to decrease the mouse’s excitability of a novel environment. After, the microscope was powered on and recording sessions were held for 5 or 20 minutes. For the 5-minute session dataset, the 5-minute sessions were repeated for 3 consecutive days with each excitation light value.

### GEVI In vivo head-fixed imaging

Surgical procedures were described in detail by Taxidis et al. 2020 [33]. Briefly, mice were anaesthetized with isoflurane and stereotactically injected with 500nl undiluted AAV8-ef1α-DiO-ASAP3-Kv (titer 3.83×1012 µg/ml), using a Nanoject II microinjector (Drummond Scientific). The injection location was in the right dorsal CA1 area (−2mm posterior and 1.8 mm lateral to bregma, 1.3 mm ventral from dura) at 60 nl/min. 1 hour later, a 3mm diameter craniotomy was performed around the injection site and the tissue above the dorsal CA1 region was aspirated. Then, a 3-mm titanium ring with a glass coverslip attached to its bottom was implanted above the CA1 region and glued to the skull. Also, a custom lightweight metal head holder (headbar) was attached to the skull posterior to the implant. Cyanoacrylate glue and black dental cement (Ortho-Jet, Lang Dental) were then added to seal and cover the exposed skull. Post-surgery recovery medications of the mice were carprofen (5 mg per kg of body weight) for 3 days post-surgery, as well as amoxicillin antibiotic (0.25 mg ml−1 in drinking water) through the water supply for 5 days.

After 4 weeks of post surgery recovery and 2 days of habituation to head fixation, mice were positioned on a custom running wheel where a custom-built, single-photon, epi-fluorescent bench microscope was used for the initial GEVI previously described in Taxidis et al. 2025 [34]. The microscope setup uses a fiber-coupled LED (Thorlabs, M455F3) with a center wavelength of 455nm where excitation light was collimated after a 2-m long, 400-μm core multi-mode fiber optic patch cord (Thorlabs, M28L02) and expanded using a Keplerian telescope. The resulting beam is then received by a spectral excitation filter (Thorlabs, MF455-45), and reflected off of a long-pass dichroic mirror (Thorlabs, MD480) before reaching a 16×/0.8NA water-immersion objective (Nikon, CFI75 LWD 16X W). This setup creates a localized excitatory spot ∼165 μm in diameter, at the focal plane. The emitted fluorescence is passed through a dichroic mirror and an emission filter (Thorlabs, MF530-43) before reaching a 100-mm tube lens (Thorlabs, AC300-100-A) followed by the imaging camera. A development board (IDS U3-3862LE-M) with a Sony IMX290LLR image sensor was used as the imaging camera. The IDS development board comes with IDS custom GUI for camera operation and image capture.

After GEVI imaging verification of the mice, the custom bench microscope was removed and MiniFAST was secured to a bar controlled by a micro-manipulator. The MiniFAST setup was placed over the same mouse/running wheel and the position of MiniFAST was finely controlled by the micro-manipulator.

### Image analysis

#### High resolution test target analysis

To calculate the contrast in the high-resolution test target images, using MATLAB, a rectangular region of interest (ROI) was manually selected over the line pairs for each group and element numbers. For each ROI, a line profile was calculated by averaging the pixel intensities across every row. Then, the contrast of each element number was calculated as (I_max_-I_min_)/(I_max_ + I_min_) where I_max_ is the peak intensity and I_min_ is the

#### *In vitro* imaging analysis

To extract the fluorescence voltage activity of the HEKS cells expressing ACE2N-mNeon, using MATLAB, an ROI was manually selected around the cell soma in one frame. Then, to find the fluorescence intensity, *F(t)*, the ROI was averaged for each frame. For each pulse trial, the baseline F was the average of *F(t)* across a 100-ms time when no voltage pulses were applied. Then, *ΔF/F = (F(t) – F)/F*. The pulse train timestamps were aligned to the fluorescent traces.

#### Calcium traces analysis

For ROI segmentation, ImageJ was used to produce background subtracted maximum projection images which were then used in MATLAB for manual curation of ROIs. To calculate the fluorescence activity in each ROI, *ΔF/F*, MATLAB was used with the following steps. First, to filter out background noise fluctuations, the following formula was applied

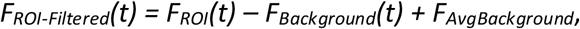

where *F*_*ROI*_*(t)* is the ROI average for each frame, *F*_*Background*_*(t)* is the entire FOV average for each frame, and *F*_*AvgBackground*_ is the average of *F*_*Background*_*(t)* across all frames. Then,

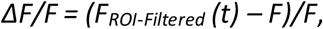

where *F* is the average of *F*_*ROI-Filtered*_*(t)* across all frames.

#### *in vivo* GEVI image analysis

For analysis of the neural data from the *in vivo* head-fixed GEVI imaging, NOSA^32^, an analytical toolbox for multicellular optical electrophysiology, was used for the data sets from both the IMX290 development board and the MiniFAST imaging. NOSA, was designed specifically for the analysis of GEVIs for the unique challenges that GEVI imaging presents. Each recording was loaded into the software, an ROI was placed over the cell, a second was placed for the background and a third ROI was placed for an off-cell analysis off the main cell, but nearby. The settings for each cell recording were the following: Background subtraction - ROI, Baseline filtering – Moving average with window 100, Spike detection – Dynamic threshold – smooth and relative threshold – 4.5 std of noise. As shown in Supplementary Figure 9, the same settings were used across each recording, for on cell, off-cell and the on-cell inverted. The data was then exported (NOSA generates an Excel sheet of the data samples, spikes, etc) into Matlab and timestamped using the MiniFAST GUI timestamp file generated during recording, for exact timing to be displayed on the figure.

#### Photobleaching analysis

To calculate the rate of photobleaching for the five-minute sessions, ImageJ was used to produce a stack of maximum projections images for every 1000 frames. Then, using MATLAB, an ROI was selected encompassing the majority of the FOV but excluding regions where the microscope housing was visible. Then, for each five-minute session, the initial fluorescence was calculated as the ROI average over the first 4 maximum projection images (∼1 minute) and the final fluorescence was calculated as the ROI average over the last 4 maximum projection images (∼1 minute). MATLAB’s Statistics and Machine Learning Toolbox was used to perform the ANOVA 2-way test with input factors of the 2 excitation light powers and the 4 mice’s 3 separate trials. For the 20-minute session, the ROI average was calculated using a 3-minute moving window and plotted.

#### Place field analysis

Place fields were computed from an approximately 17-minute-long session (Fig. 2d, 2e, Supplementary Fig. 3). Videos were first motion corrected with CaImAn^29^. Next, the maximum projection image computed over the videos was used to obtain initial ROIs from an automatic ROI detection algorithm in CaImAn (Fig. 2b). These ROIs were used to initialize the spatial footprints for CNMF-E and to plot fluorescence traces (Fig. 2c). As CA1 activity is sparse, for simplicity of display example fluorescence traces use the mean over an ROI for the baseline.

Following CNMF-E deconvolution, the vector of spiking activity was extracted for each ROI. Since these vectors were non-integral valued, they were treated as pseudo spike counts. Animal position was determined by tracking a LED on the microscope assembly. For each ROI, a ratemap was generated by binning the spiking activity vector into 2D spatial bins of dimensions 1 cm × 1 cm and normalizing by the occupancy. Only periods of time when the animal was running (peak speed > 10 cm/s with a threshold of 8 cm/s as the running event boundaries) were considered. The ratemaps were smoothed with a Gaussian kernel of standard deviation 2 cm and normalized to the range [0, 1] for visualization (Fig. 2d, 2e and Supplementary Fig. 3).

For each occupancy-normalized ratemap, “nelpy” a custom Python package https://github.com/nelpy/nelpy, was used to compute the spatial information^30^. Next the position timestamps were circularly shifted 1000 times by a random amount to generate a shuffle distribution of spatial information. A ratemap was considered a valid place field if it satisfied the following criteria: (1) its corresponding spiking activity vector was nonzero at least 5 times, (2) the total number of non-zero elements in the spiking activity vector divided by the session duration resulted in an overall frequency of < 0.5 Hz, and (3) the spatial information was greater than the 95^th^ percentile of the shuffle distribution.

#### Project design availability

The MiniFAST 3D-printed housing, hardware, DAQ firmware and updated Miniscope GUI is available at https://github.com/jjuneau1/MiniFAST. Additionally, in the same repository, there are 3-D printed design files for a 1.8mm GRIN lens surgical holder and the MiniFAST patch clamp setup brackets.

## Availability of data and analysis code

The experimental data collected and analyzed in this publication is available from the corresponding author upon reasonable request.

The custom analysis code in this publication is available from the corresponding author upon reasonable request.

## Acknowledgements

We thank the Miniscope Team, Framos Technology, the Rice University ARF, E. Orchard, T. Volz, E. Festa and S. Gao for conversations and assistance. This work was supported by funding from the NIH (R21EY029459 and RF1NS110501) and DARPA (N66001-17-C-4012).

## Competing Interests

The authors declare no competing interests.

## Notes

### Competing Interest Statement

The authors have declared no competing interest.

### Summary of Updates

This version of the manuscript has been revised to include new results from experiments using MiniFAST with GEVI in vivo head-fixed mice.

