## Supplementary Figures for "MiniFAST: A sensitive and fast miniaturized microscope for *in vivo* neural recording"

### Supplementary Materials

#### Supplementary Table 1: Currently available miniaturized microscopes published specifications

| Microscope Name | Frame Rate | Neural Indicator (animal) | Sensor Resolution | Pixel Size |
|---|---|---|---|---|
| UCLA Miniscope V3.2 [3] | 60 Hz | AAV–GCaMP6f (mouse) | 752x480 pixels | 6um |
| miniScope [20] | 10 Hz | AAV-GCaMP6s (mouse) | 400x400 pixels | 6um |
| Wirefree miniScope[6] | 10 Hz | AAV-GCaMP6s (mouse) | 200x200 pixels | 6um |
| Finchscope[4] | 30 Hz | AAV-GCaMP6s and GCaMP6f (bird) | 640x480 pixels | Not published |
| cScope [7] | 60 Hz | Transgenic Thy1-GCAMP6f (rat) | 752x480 pixels | 6um |
| Multi-Contrast[21] | 15 Hz | Transgenic GCaMP6s (mouse) | 640x640 pixels | 3um |
| MiniLFM [9] | 16 Hz | AAV-GCaMP6f (mouse) | 1280x1024 pixels | 5.2um |
| NINscope [8] | 30 Hz | AAV-GCaMP6f (mouse) | 808x608 pixels | 4.8um |
| Dual Hemisphere [10] | 25 Hz | Transgenic Thy1-GCaMP6s (mouse) | 720x576 pixels | 6um |
| Wirefree UCLA Miniscope[5] | 20 Hz | AAV-GCaMP6f (mouse) | 836x640 pixels | 5.8um |

#### Supplementary Figure 1: MiniFAST image sensor PCB overview

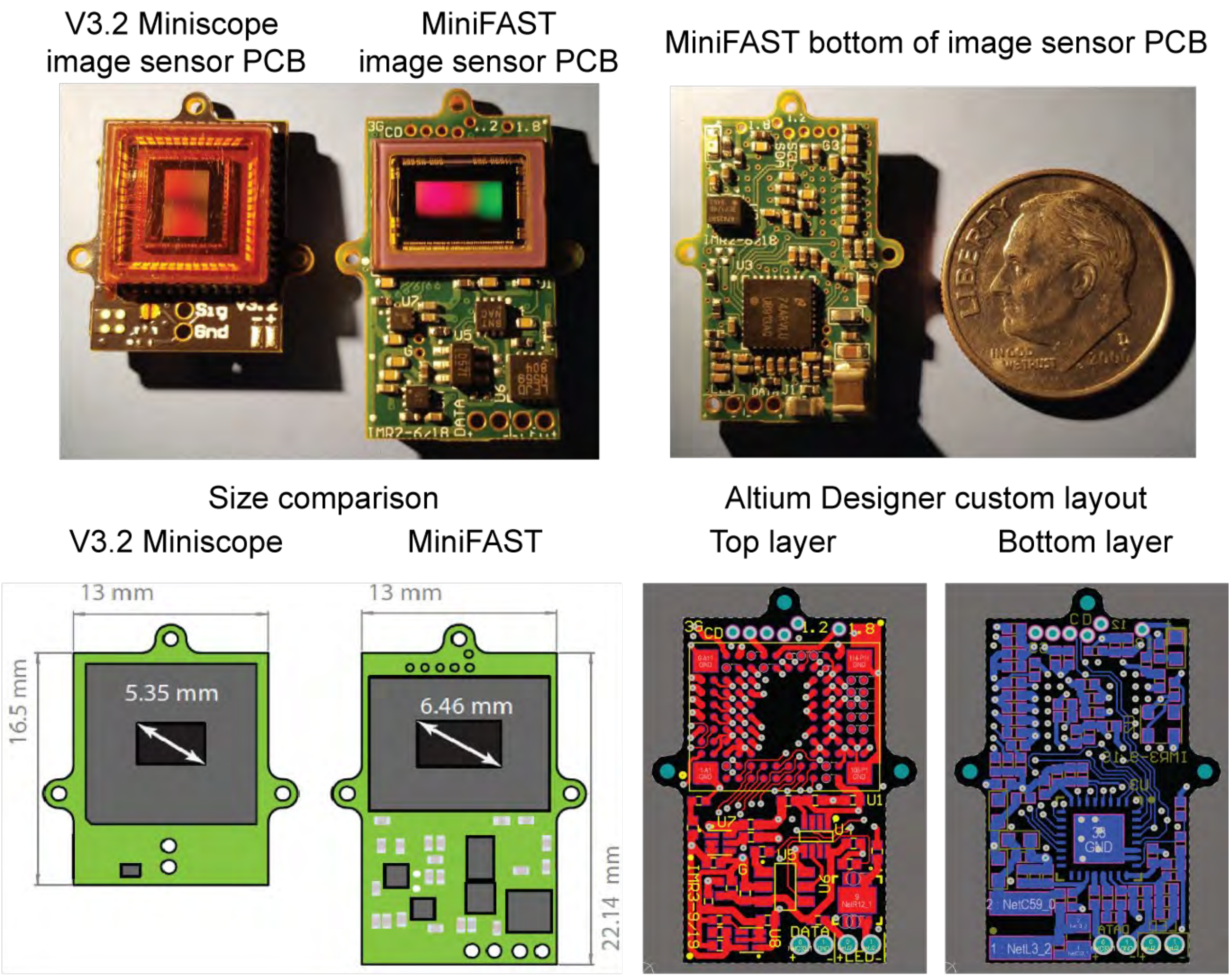

**Adapted from figure in (Juneau et al., 2018[29])**

#### Supplementary Table 2: Miniaturized microscope image sensor specification comparison

| | Miniaturized Microscopes | | |
|---|---|---|---|
| **Parameters** | **V3.2 UCLA Miniscope** | **NINScope** | **MiniFAST** |
| **Weight (PCB,housing, optics)** | 3.2 g | 1.7 g | 3.45 g |
| **Image Sensor Manufacturer** | On Semi | On Semi | Sony |
| **Image Sensor Model** | MT9V032 | Python 480 | IMX290LLR-C |
| **Shutter** | Global | Global | Rolling |
| **Resolution (pixels)** | 752 x 480 | 808 x 608 | **1920 x 1080 @ 30 Hz** |
| **Pixel Size** | 6 um | 4.8 um | **2.9 um** |
| **Sensor Format** | 1/3" | 1/3.6" | 1/2.8" |
| **Maximum Gain** | 12 dB | 11 dB | **30 dB (analog)<br>+ 42 dB (digital)** |
| **Max Frame Rate** | 60 Hz | 120 Hz<br>(NINscope @ 30Hz) | **> 500 Hz<br>(with decreasing window size)** |

### Supplementary Figure 2: High resolution test target extended figures

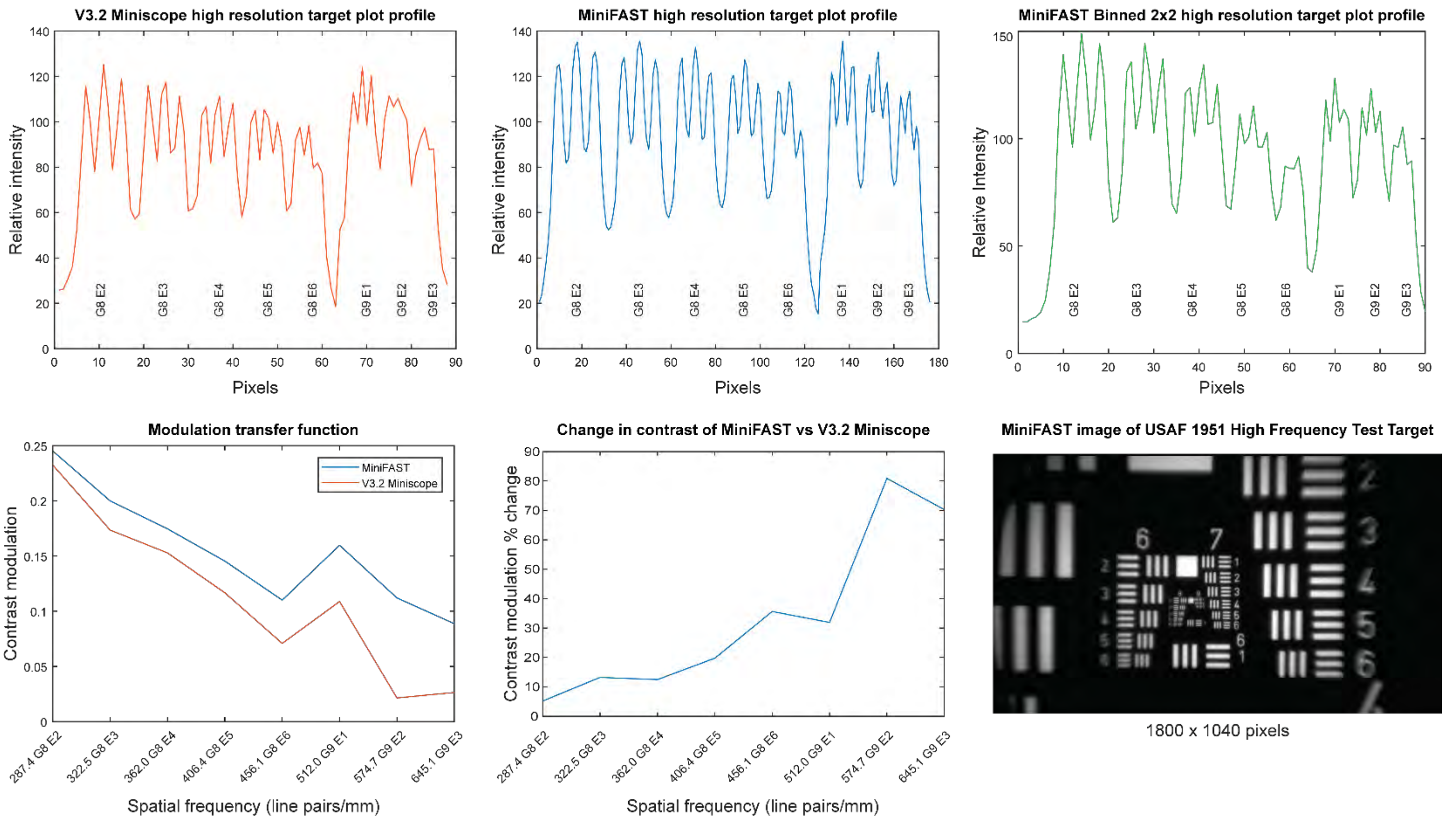

#### Supplementary Figure 3: Detected place fields from exploration session

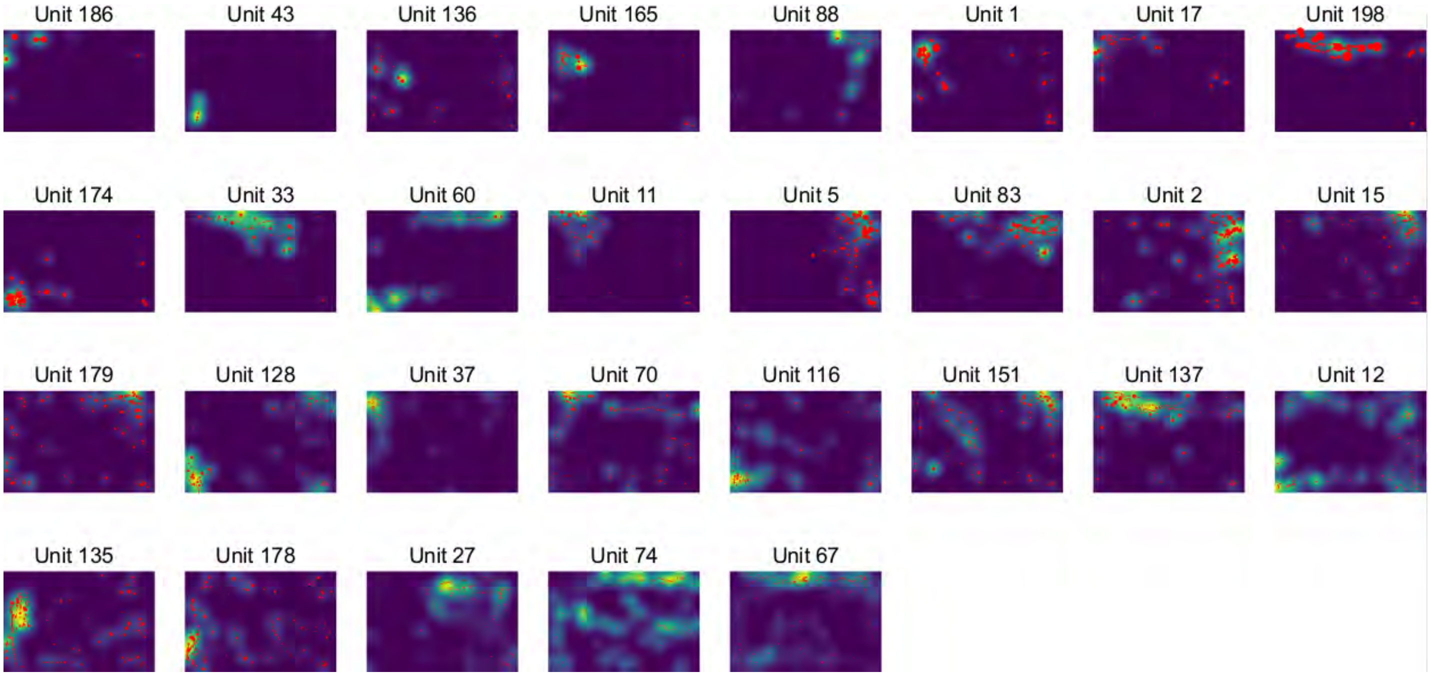

Deconvolved spikes (red) overlaid on inferred ratemaps for 29 neurons which displayed fields which were more spatially tuned than chance (i.e., place cells).

#### Supplementary Figure 4: 2-Hour continuous imaging session of calcium imaging

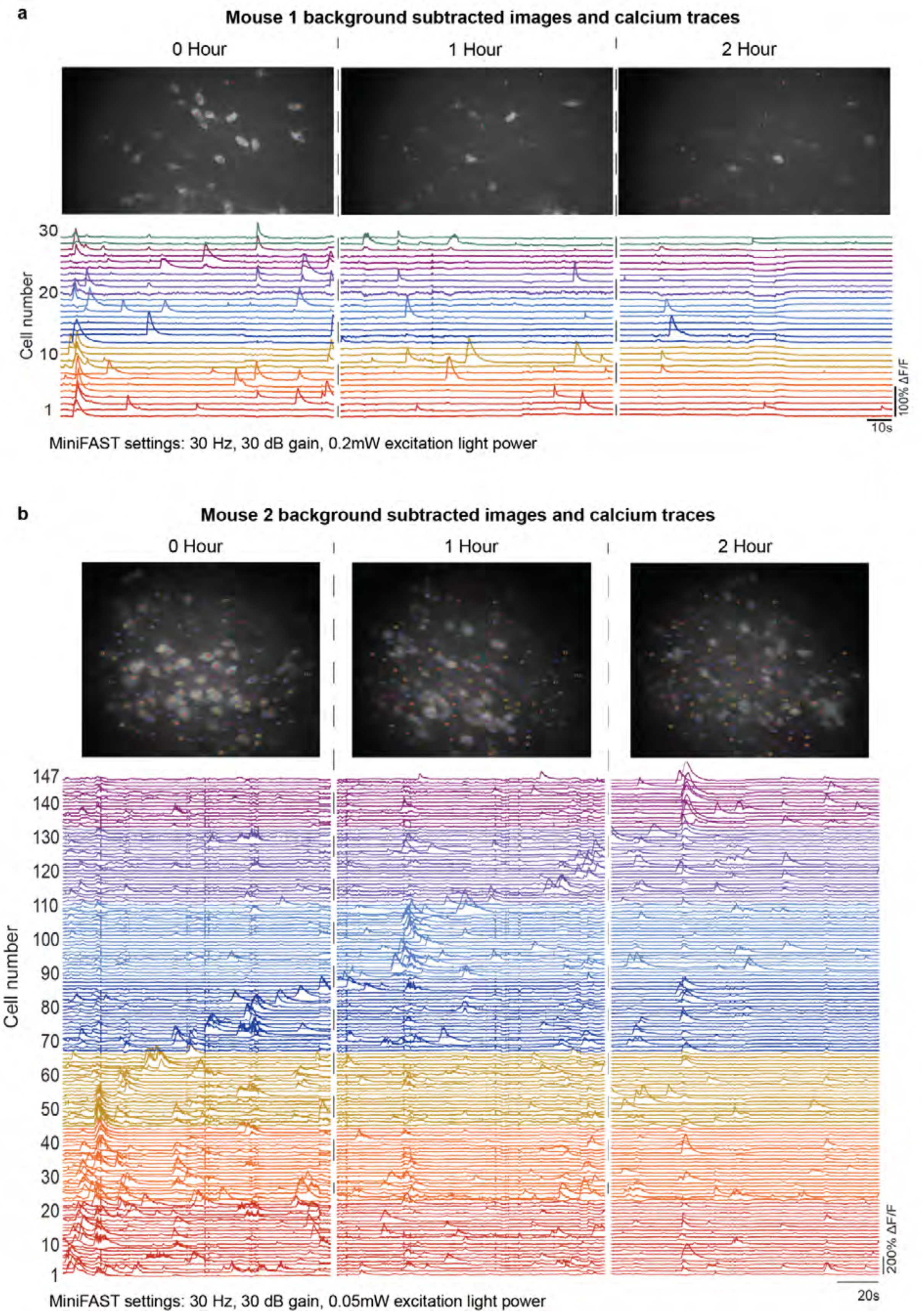

#### Supplementary Figure 5: Long-term (1-month) calcium imaging in CA1 of AAV-GCaMP6f mice

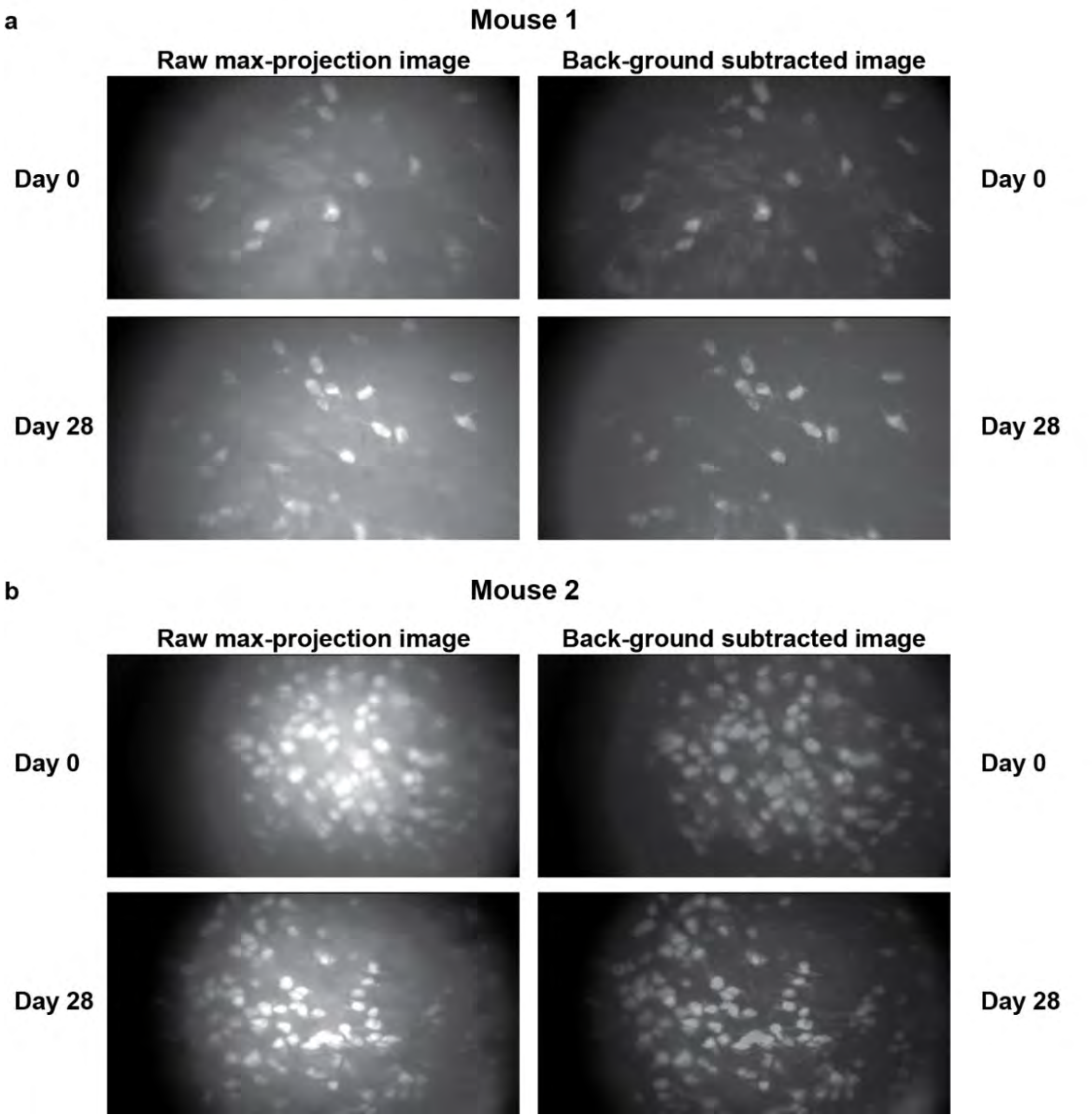

**Supplementary Figure 6: Long-term (4-month) calcium imaging in CA1 of transgenic Thy1-GCaMP6f GP5.17 mice**

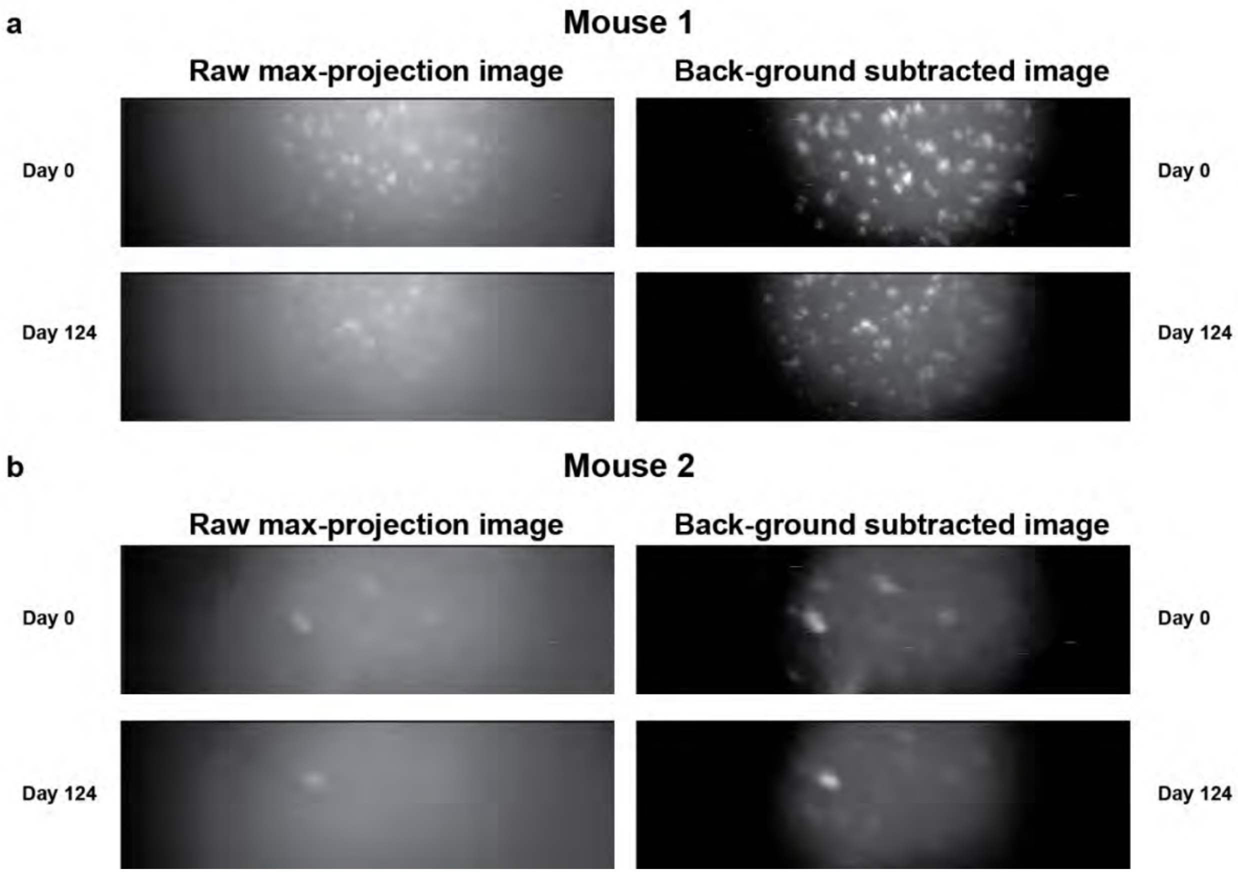

**Supplementary Figure 7: High speed imaging of CA1 of transgenic Thy1-GCaMP6f GP5.17 mice**

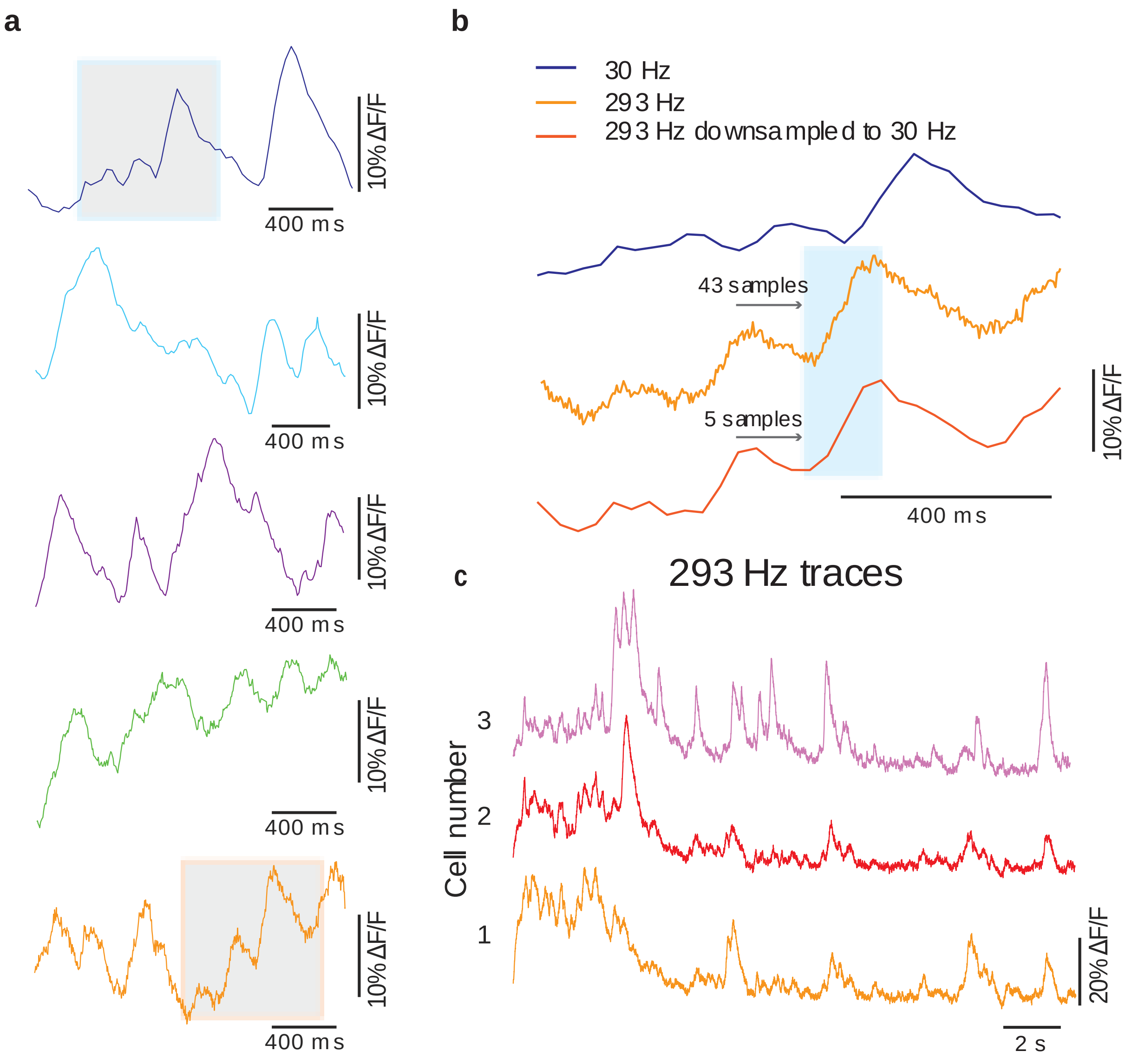

*a, 2-second zoomed in traces from (Figure 3 -**b)*** boxed sections. ***b***, 1-second zoomed traces of 30 Hz and 293 Hz from (***a***). ***c***, *$Ca^{2+}$ traces of 3 cells in the FOV at 293 Hz.* ***a***, *Settings: 30 Hz (1800 x 1040 pixels), 27 dB gain, 0.86 mW excitation light power.* ***b***, *Settings: 60 Hz (1900 x 544 pixels), 27 dB gain, 1.4 mW excitation light power.* ***c***, *Settings: 100 Hz (1900 x 337 pixels), 27 dB gain, 2.5 mW excitation light power.* ***d***, *Settings: 200 Hz (1900 x 156 pixels), 30 dB gain, 2.5 mW excitation light power.* ***e***, *Settings: 293 Hz (1900 x 102 pixels), 30 dB gain, 3.25 mW excitation light power.*

#### Supplementary Figure 8: Photobleaching evaluation of CA1 of transgenic Thy1-GCaMP6f GP5.17 mice

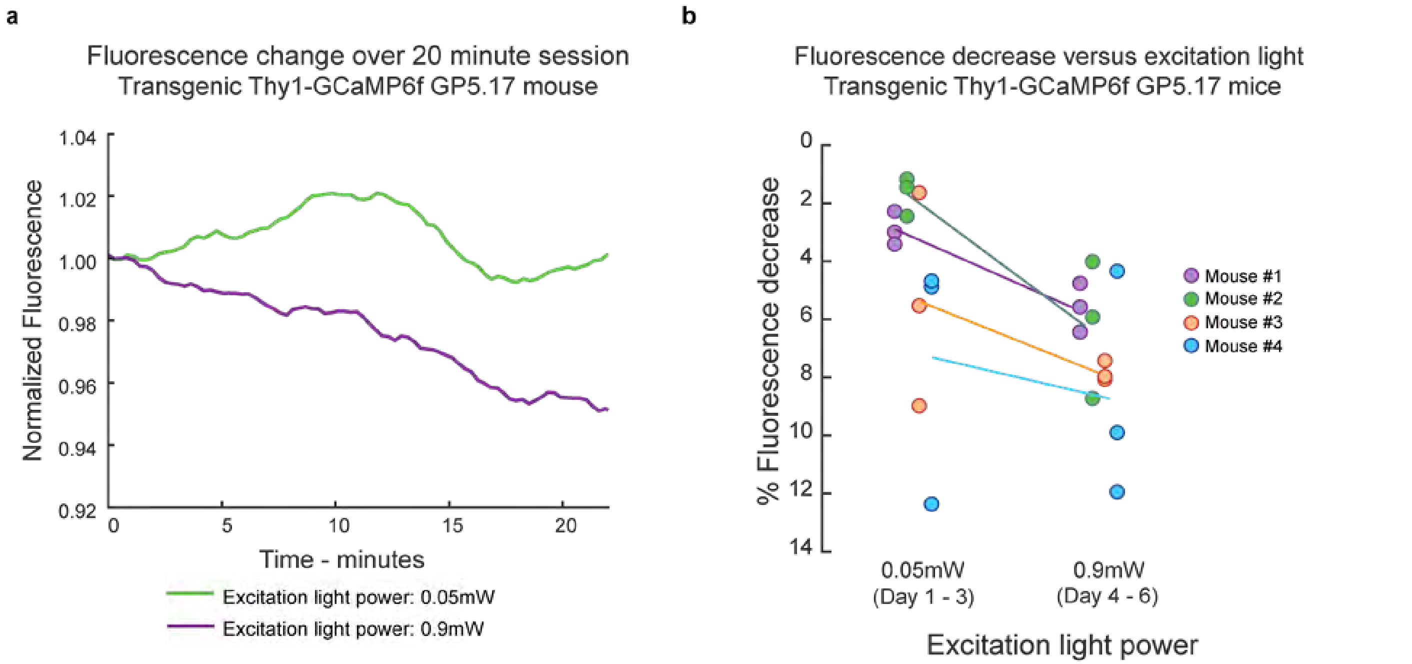

***a**, Plot of fluorescence change over a 20-minute imaging session of a Thy1-GCaMP6f GP5.17 mouse with 2 different excitation light power settings. **b**, Scatter plot of fluorescence decrease versus excitation light power for 4 Thy1-GCaMP6f GP5.17 mice during 5-minute imaging sessions occurring over 6 days. For each mouse, one imaging session occurred per day and days are noted on plot below excitation light power. Each circle represents one session and each color represents a different mouse. Settings for 0.05 mW excitation power, 60 Hz, 51 dB gain. Settings for 0.9mW excitation light power, 60 Hz, 27 dB gain.*

Since MiniFAST has the ability to image at fast frame rates in low light conditions, we wanted to evaluate whether imaging with high gain settings and low excitation light values could help reduce photobleaching effects while still being able to track $Ca^{2+}$ activity at frame rates of 60 Hz. To understand how the excitation light power influences photobleaching, we continuously imaged a Thy1-GCaMP6f GP5.17 mouse over two 20-minute open field sessions where one session used a very low excitation light power of 0.05 mW and the other session used a higher excitation light power of 0.9 mW (Fig. 5a). Plotting the average fluorescence over the image FOV, the normalized fluorescence was impacted minimally for the extremely low power setting of 0.05 mW and decreased by ~5% for the higher excitation light power of 0.9 mW. To examine this effect in depth, over a total period of 6 days, we performed 5-minute imaging sessions each day (n = 4 mice) with sessions of 0.05 mW and 0.9 mW excitation light power settings (Fig. 5b). For each session, the difference of the average fluorescence of the first and last minute was calculated. Although there was some variation in the day-to-day changes, higher excitation light power generally resulted in a significantly greater decrease in the fluorescence signal ($p < 0.05$ ANOVA, $F(1, 3) = 6.7$, $p = 0.02$).

### Supplementary Figure 9: NOSA GUI interface

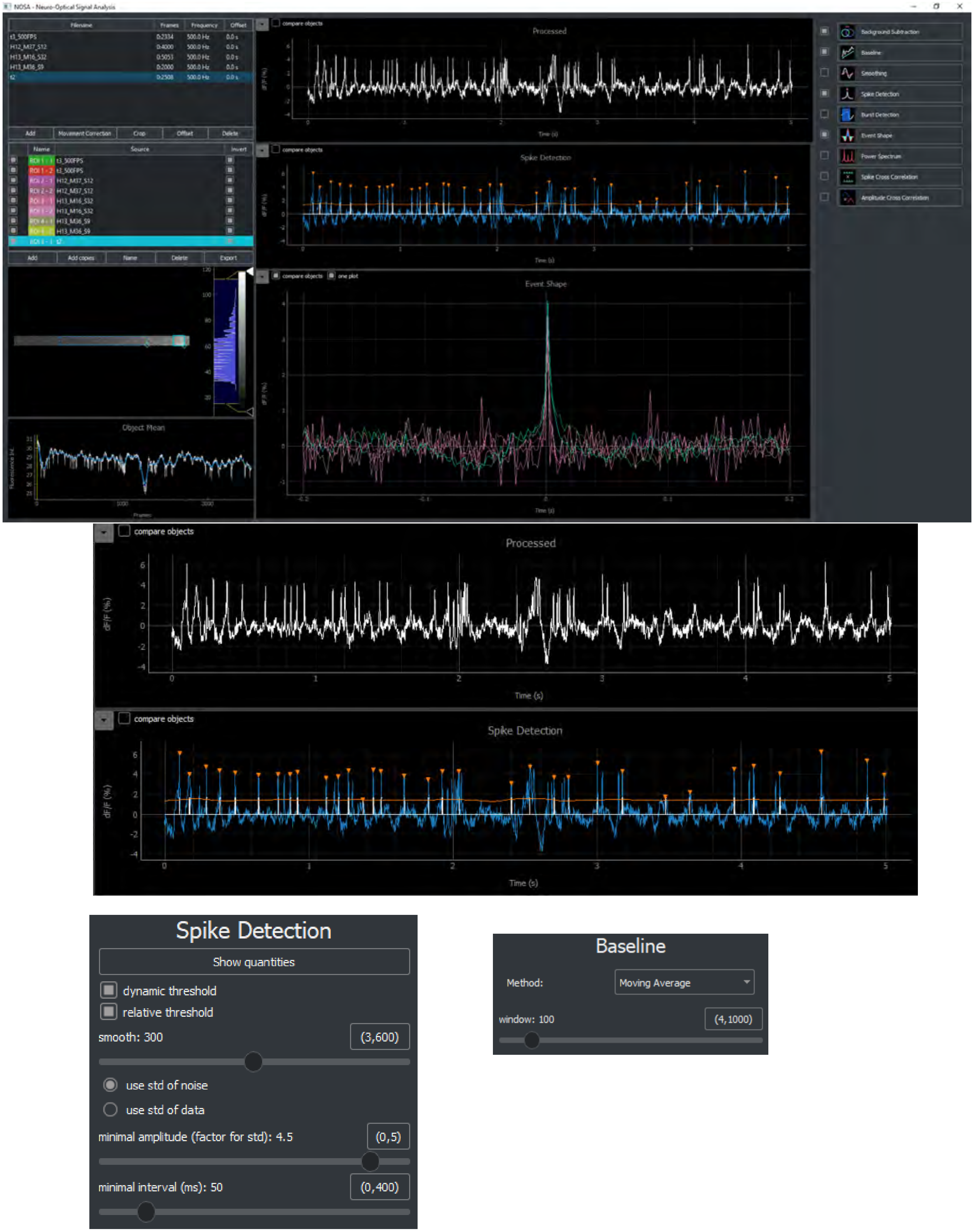

**Supplementary Figure 10: MiniFAST *in vivo* imaging of ASAP GEVIs in hippocampal SST $^{+}$ interneurons in head-fixed mice – Cell 1 detail**

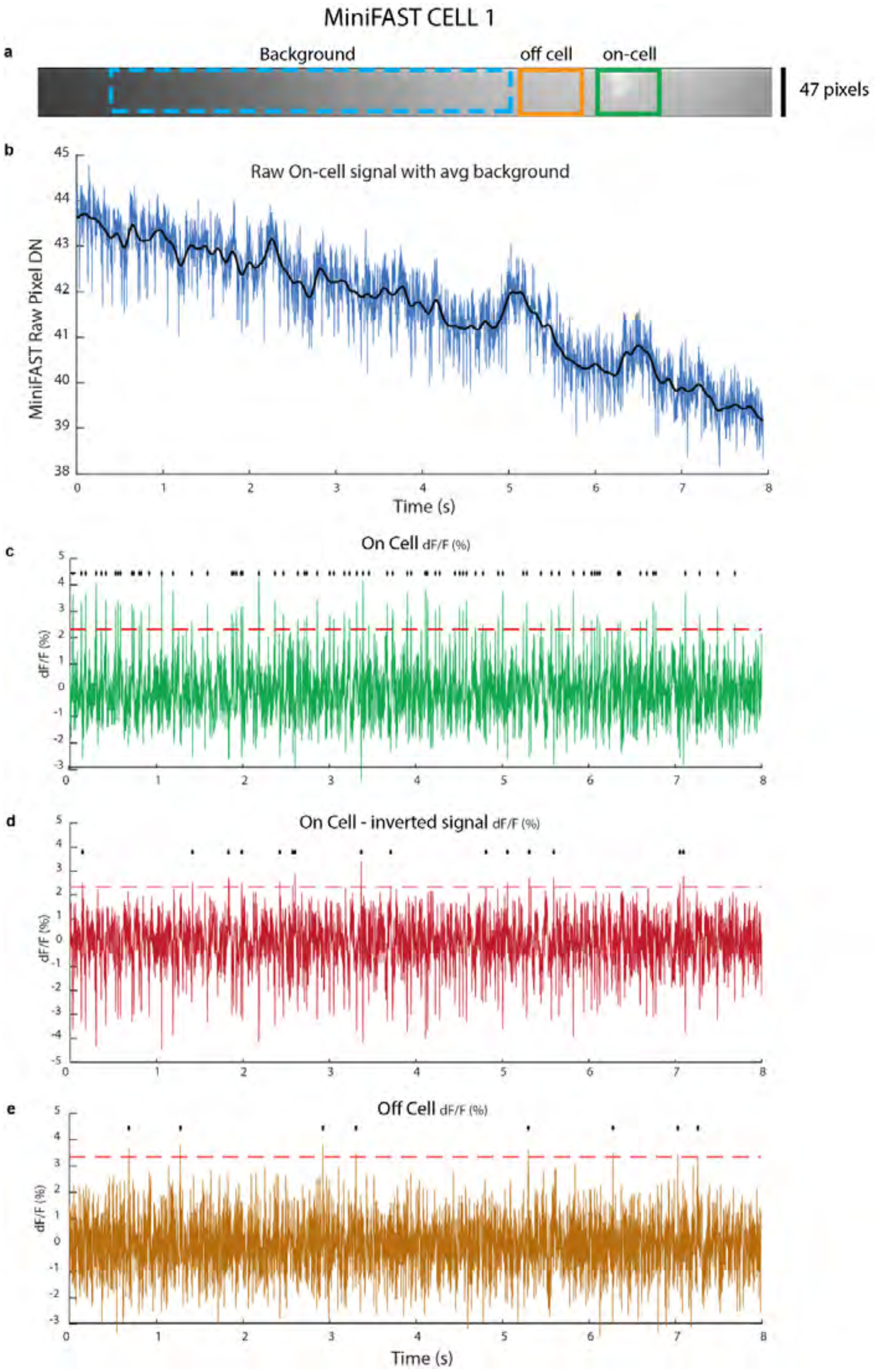

***a**, ImageJ z-projection standard deviation image of MiniFAST ASAP3-Kv cell with highlighted boxes for the cell ROI, an off-cell region and background area. **b**, Raw GEVI trace (not-inverted) with pixel DN counts – MiniFAST output is 8-bits. ASAP3-Kv fluorescent traces were acquired at 500fps, camera gain of 37db, using light excitation power of 10mW mm$^{-2}$. **c,** On-cell GEVI trace of background subtracted signal. ASAP3-Kv is a high to low indicator and the traces shown here have been inverted. The red dashed line represents a spike threshold of 4.5 standard deviations of the noise and black dots represent detected spikes. **d**, Same trace as **(c)**, but inverted and reprocessed with red line representing 4.5 standard deviations of the noise. **e**, Off-cell ROI signal trace with red line representing 4.5 standard deviations of the noise.*

**Supplementary Figure 11: MiniFAST *in vivo* imaging of ASAP GEVIs in hippocampal SST$^{+}$ interneurons in head-fixed mice – Cell 2 detail**

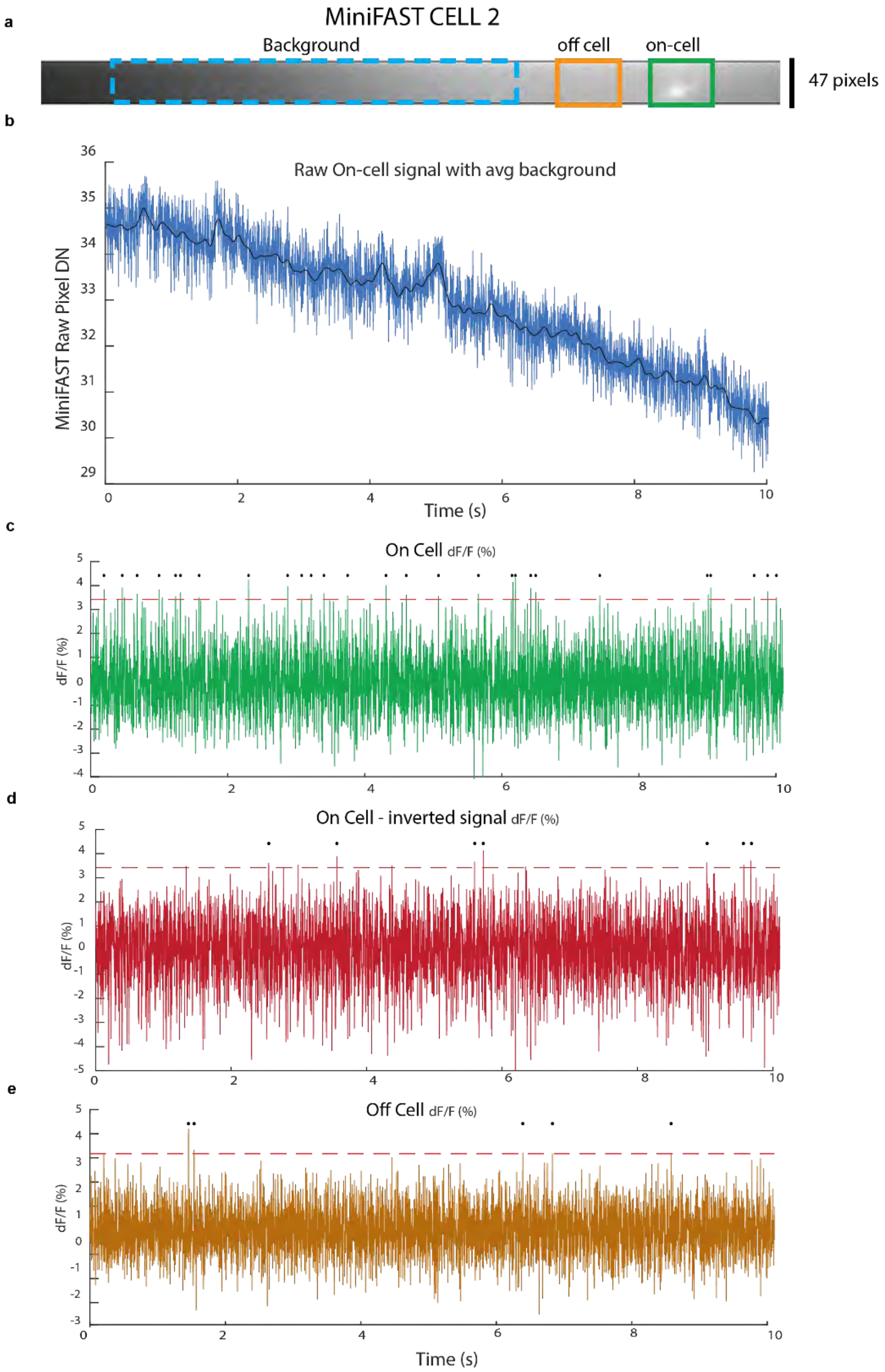

***a***, *ImageJ z-projection standard deviation image of MiniFAST ASAP3-Kv cell with highlighted boxes for the cell ROI, an off-cell region and background area.* ***b***, *Raw GEVI trace (not-inverted) with pixel DN counts – MiniFAST output is 8-bits. ASAP3-Kv fluorescent traces were acquired at 500fps, camera gain of 37db, using light excitation power of 10mW mm$^{-2}$.* ***c,*** *On-cell GEVI trace of background subtracted signal. ASAP3-Kv is a high to low indicator and the traces shown here have been inverted. The red dashed line represents a spike threshold of 4.5 standard deviations of the noise and black dots represent detected spikes.* ***d***, *Same trace as* ***(c)***, *but inverted and reprocessed with red line representing 4.5 standard deviations of the noise.* ***e***, *Off-cell ROI signal trace with red line representing 4.5 standard deviations of the noise.*

**Supplementary Figure 12: MiniFAST *in vivo* imaging of ASAP GEVIs in hippocampal SST $^{+}$ interneurons in head-fixed mice – Cell 3 detail**

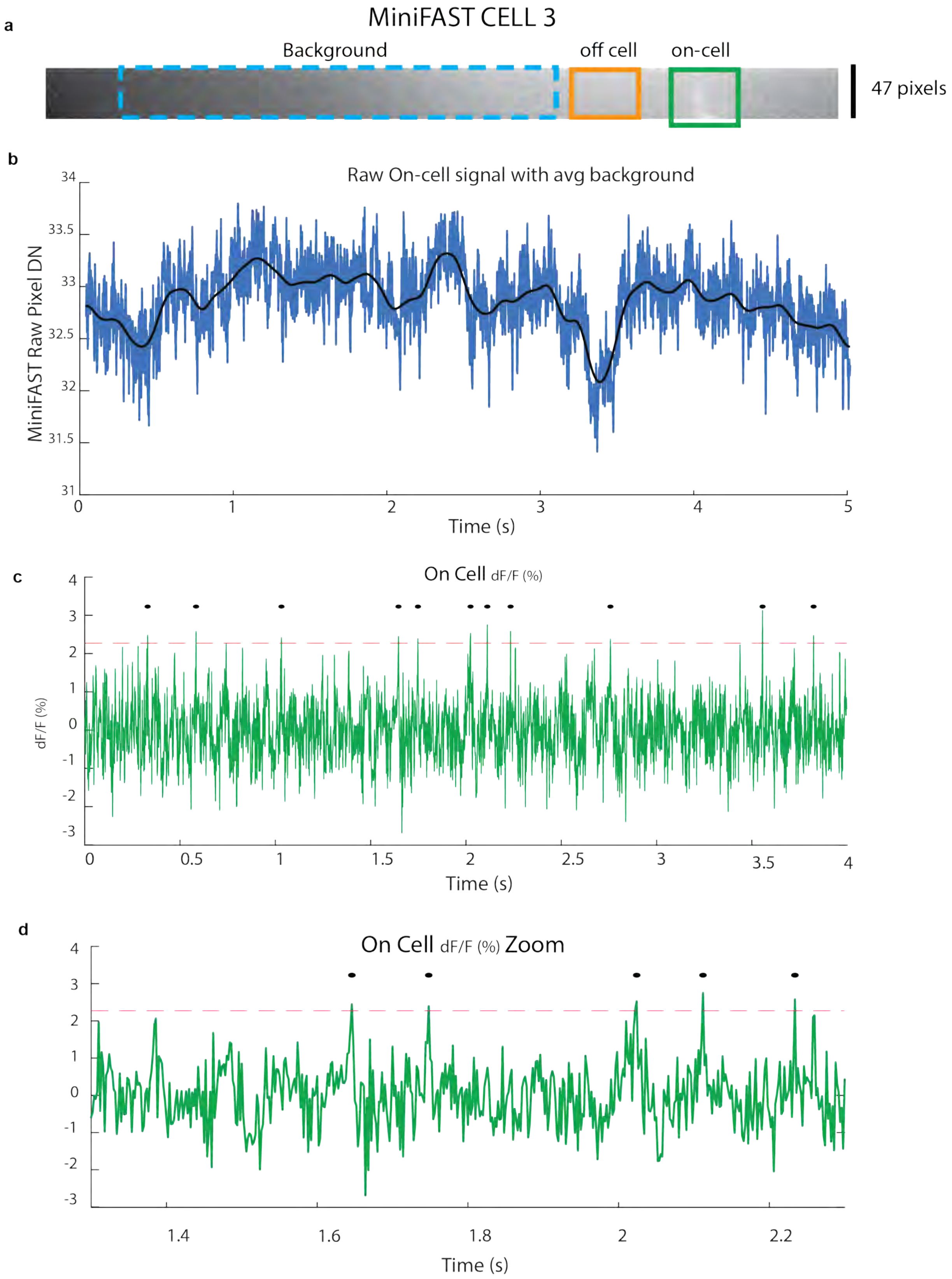

#### MiniFAST CELL 3

**e**

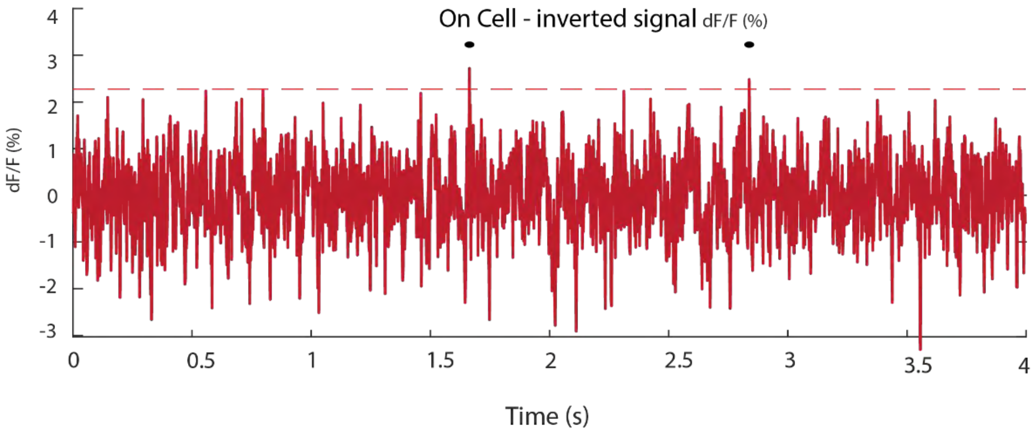

**f**

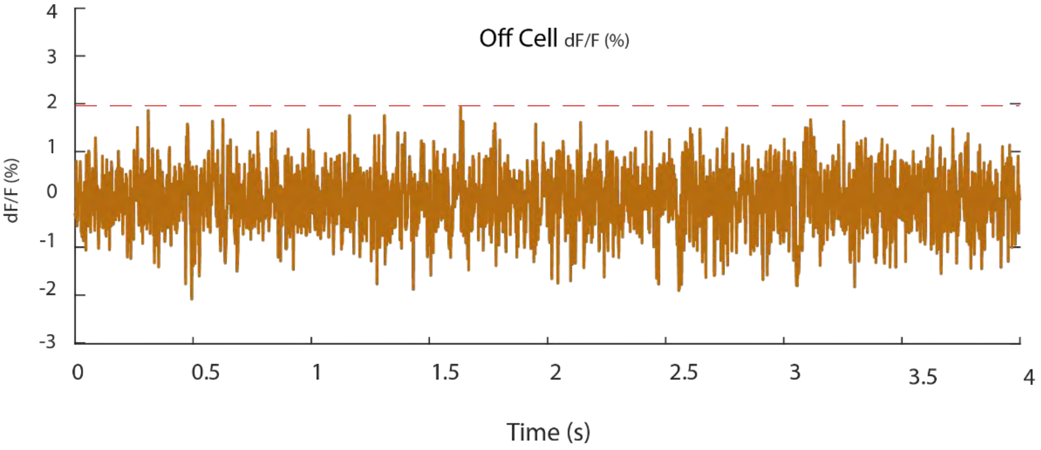

***a**, ImageJ z-projection standard deviation image of MiniFAST ASAP3-Kv cell with highlighted boxes for the cell ROI, an off-cell region and background area. **b**, Raw GEVI trace (not-inverted) with pixel DN counts – MiniFAST output is 8-bits. ASAP3-Kv fluorescent traces were acquired at 500fps, camera gain of 37db, using light excitation power of 10mW mm$^{-2}$. **c,** On-cell GEVI trace of background subtracted signal. ASAP3-Kv is a high to low indicator and the traces shown here have been inverted. The red dashed line represents a spike threshold of 4.5 standard deviations of the noise and black dots represent detected spikes. **c,** Zoomed On-cell GEVI trace of background subtracted signal. **e**, Same trace as **(c)**, but inverted and reprocessed with red line representing 4.5 standard deviations of the noise. f, Off-cell ROI signal trace with red line representing 4.5 standard deviations of the noise.*

**Supplementary Figure 13: MiniFAST *in vivo* imaging of ASAP GEVIs in hippocampal SST + interneurons in head-fixed mice – Average spike trace from all cells**

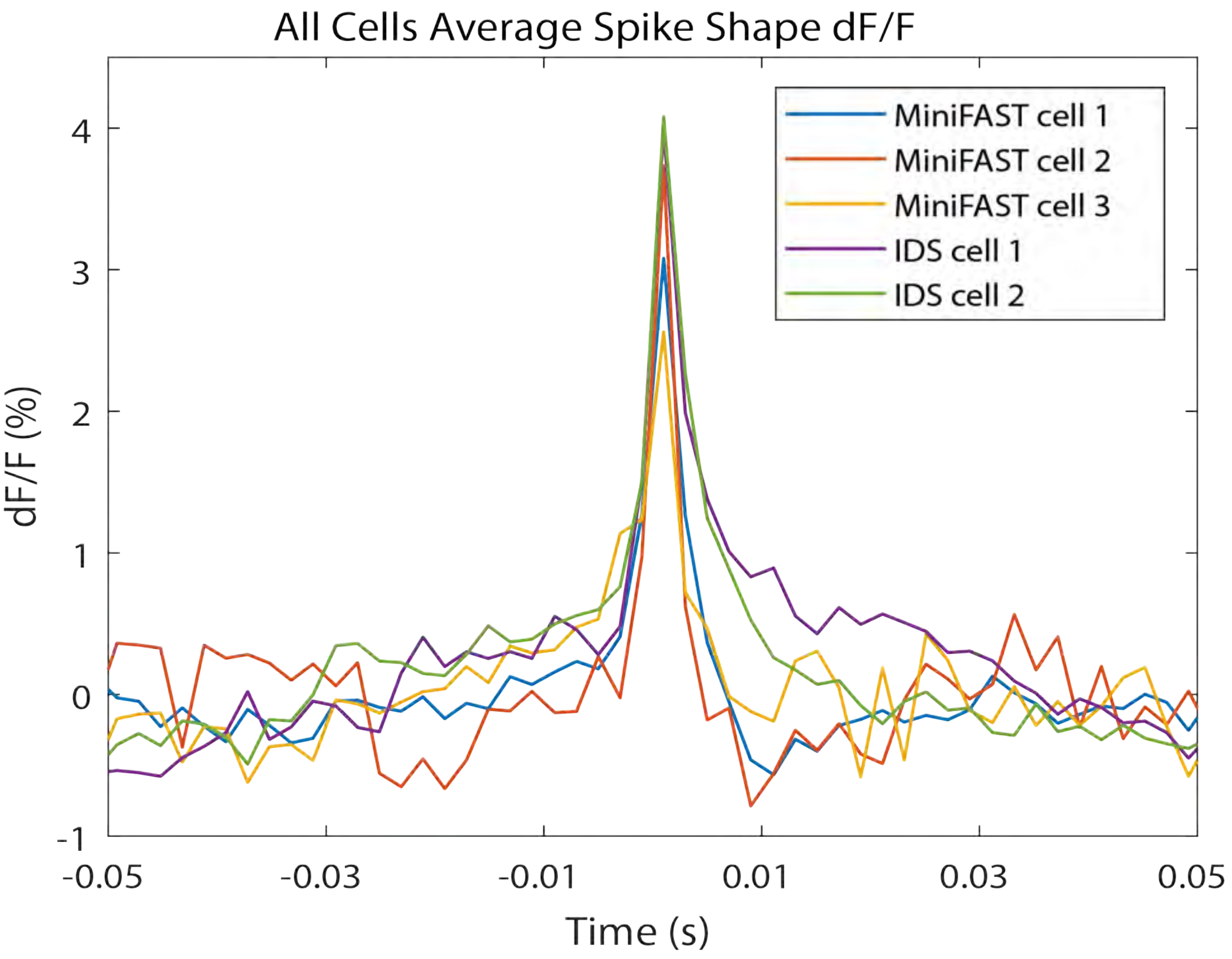

*Average spike event shape for the 2 cells shown for the IMX290 development board setup and the 3 cells from the MiniFAST setup.*

**Supplementary Figure 14: Sony IMX290 row size vs frame rate for the parallel/CSI interfaces**

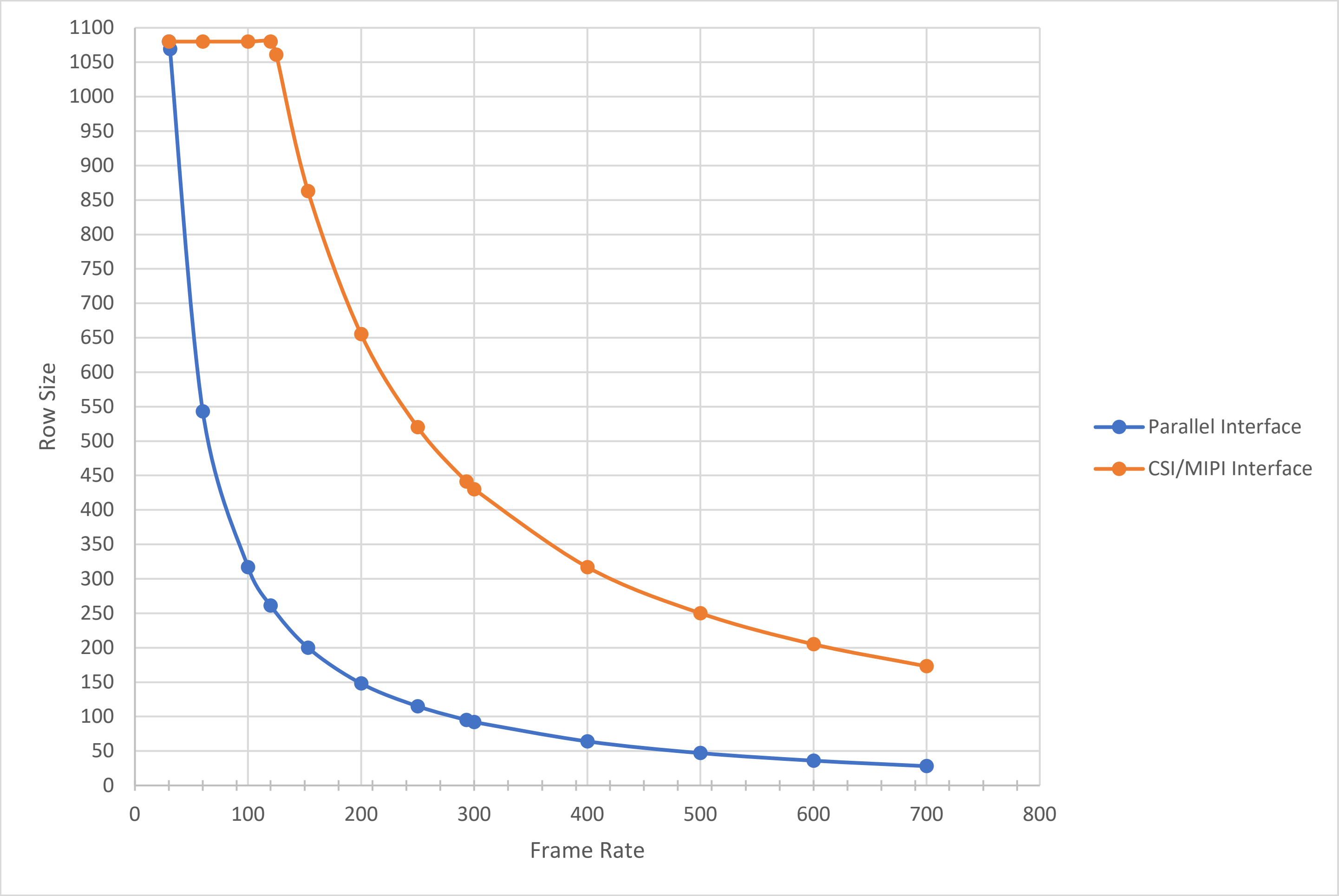
